# PHASE: A MATLAB Based Program for the Analysis of *Drosophila* Phase, Activity and Sleep under Entrainment

**DOI:** 10.1101/2021.12.14.472617

**Authors:** J.L. Persons, L. Abhilash, A.J. Lopatkin, A. Roelofs, E.V. Bell, M.P. Fernandez, O.T. Shafer

## Abstract

The problem of entrainment is central to circadian biology. In this regard, *Drosophila* has been an important model system. Owing to the simplicity of its nervous system and the availability of powerful genetic tools, the system has shed significant light on the molecular and neural underpinnings of entrainment. However, much remains to be learned regarding the molecular and physiological mechanisms underlying this important phenomenon. Under cyclic light/dark conditions, *Drosophila melanogaster* displays crepuscular patterns of locomotor activity with one peak anticipating dawn and the other anticipating dusk. These peaks are characterized through an estimation of their phase relative to the environmental light cycle and the extent of their anticipation of light transitions. In *Drosophila* chronobiology, estimations of phases are often subjective, and anticipation indices vary significantly between studies. Though there is increasing interest in building flexible analysis software tools in the field, none incorporates objective measures of *Drosophila* activity peaks in combination with the analysis of fly activity/sleep in the same program. To this end, we have developed PHASE, a MATLAB-based program that is simple and easy to use and (i) supports the visualization and analysis of activity and sleep under entrainment, (ii) allows analysis of both activity and sleep parameters within user-defined windows within a diurnal cycle, (iii) uses a smoothing filter for the objective identification of peaks of activity (and therefore can be used to quantitatively characterize them), and (iv) offers a series of analyses for the assessment of behavioral anticipation of environmental transitions.

## Background/Introduction

Entrainment to environmental cycles is a key property of circadian clocks and is critical for maintaining the adaptive advantage of rhythmic phenomena across taxa (Pittendrigh, 1981; Vaze and Sharma, 2013). Critical markers of entrainment are (i) the stable and reproducible daily timing of behavior (phase of entrainment), (ii) systematic changes of these phases under various zeitgeber regimes, and (iii) anticipation of predictable environmental transitions (for instance, lights-ON and lights-OFF) (Pittendrigh, 1981; Moore-Ede et al., 1982; Roenneberg et al., 2005). Therefore, measures of phases and estimates of anticipation are critical to characterizing entrained behavior of organisms.

*Drosophila* has proved to be an insightful model system in which many of the molecular and neuronal underpinnings of entrainment have been revealed, owing to its ease of maintenance, its relatively simple nervous system, and the availability powerful genetic tools (Hardin, 2011; Helfrich-Förster, 2020). The daily morning and evening peaks of activity are the most prominent features of the *Drosophila* locomotor activity rhythm under light/dark (LD) cycles. The timing and amounts of activity underlying these peaks are therefore of significant interest to those using the fly to understand entrainment (Hamblen-Coyle et al., 1992; Renn et al., 1999; Grima et al., 2004; Stoleru et al., 2004). Thus, the objective estimation of the phases of these peaks and the extent of their anticipation of environmental transitions is critical for the characterization of the entrained circadian system of flies.

Recently, there has been an increasing interest in the development of open-source software and teaching tools to analyze and understand features of circadian rhythms (Schmid et al., 2011; Cichewicz and Hirsh, 2018; Geissmann et al., 2019; Abhilash and Sheeba, 2019; Cenek et al., 2020; Kostadinov et al., 2021; Hammad et al., 2021). However, to the best of our knowledge, a program that supports the standard analysis and visualization of *Drosophila* activity and sleep rhythms that is also easily accessible, incorporates objective analysis of phases of activity, and includes measures of anticipation is not currently available. To this end, we present PHASE – a freely available MATLAB program to analyze *Drosophila* <u>Ph</u>ase, <u>A</u>ctivity, and <u>S</u>leep under <u>E</u>ntrainment.

## PHASE: System Requirements, Installation, and the GUI

PHASE is a freely-available, open access, and stand-alone program written for MATLAB and hosted on GitHub (https://github.com/ajlopatkin/PHASE/tree/master/Installer%20Downloads). However, users can also run PHASE in MATLAB, which requires a current MATLAB license (Runtime R2020 or later; The MathWorks, Inc, Natick, MA). The PHASE bundle can be downloaded from GitHub (https://github.com/ajlopatkin/PHASE) and instructions in the supplemental manual can be used to install and run the program on Windows (7 or later) and Mac (10.13 or later) computers. Only data acquired using *Drosophila* Activity Monitor (DAM) systems via *DAMSystem3* and processed using the *DAMFileScan* can be used as input for PHASE (Trikinetics, Waltham, MA; https://www.trikinetics.com). The DAM system is the most commonly used apparatus for recording locomotor activity in flies, and is also widely used to measure sleep in the fly (Chiu et al., 2010; Cichewicz and Hirsh, 2018; Kostadinov et al., 2021).

The graphical user-interface (GUI) of PHASE can be divided into five sections (Fig. 1A). The first is where users choose the directory where all their files for analysis are stored (Fig. 1A-i). Input files for the analysis must be DAM-scanned channel files. The second section of the interface is “Data Settings” and is one of the most critical components of PHASE (Fig. 1A-ii). From left-to-right, “Data Settings” consists of the following: a display window which lists all the monitors that are available in the user-chosen folder (specified in the first section), a display window showing all the channels within the monitor files, a display window that shows which monitor and channels have been selected by the user for analysis, and input settings for the desired analysis (Fig. 1A-ii). Details regarding how to use these input settings are included in the figure legends and the supplemental user manual. The third section is where the user chooses which of the PHASE functions to execute (Figs. 1A-iii, B and C). The user chooses a scaling option for the *y*-axis in the plots, the folder in which to save all the plots and spreadsheets generated by PHASE, and the name of the output files in section four (Fig. 1A-iv). The buttons to execute the functions chosen in Fig. 1A-iii are in the fifth section of the user-interface (Fig. 1A-v). The three available buttons are “Sleep Analysis”, “Normalized Activity Analysis” and “Averaged Activity Analysis.” Each of these buttons will use a different data set to perform all the user-chosen analysis. In the case of “Sleep Analysis,” PHASE will use standard sleep data (see Shafer and Keene, 2021). For “Normalized Activity,” PHASE uses activity data normalized to the total activity per cycle, and for “Averaged Activity,” it uses activity data averaged over the length of the user-defined bin (see Fig. 1A-iii). We encourage users to read the figure legends and consult the user manual for detailed instructions.

**Figure 1:**
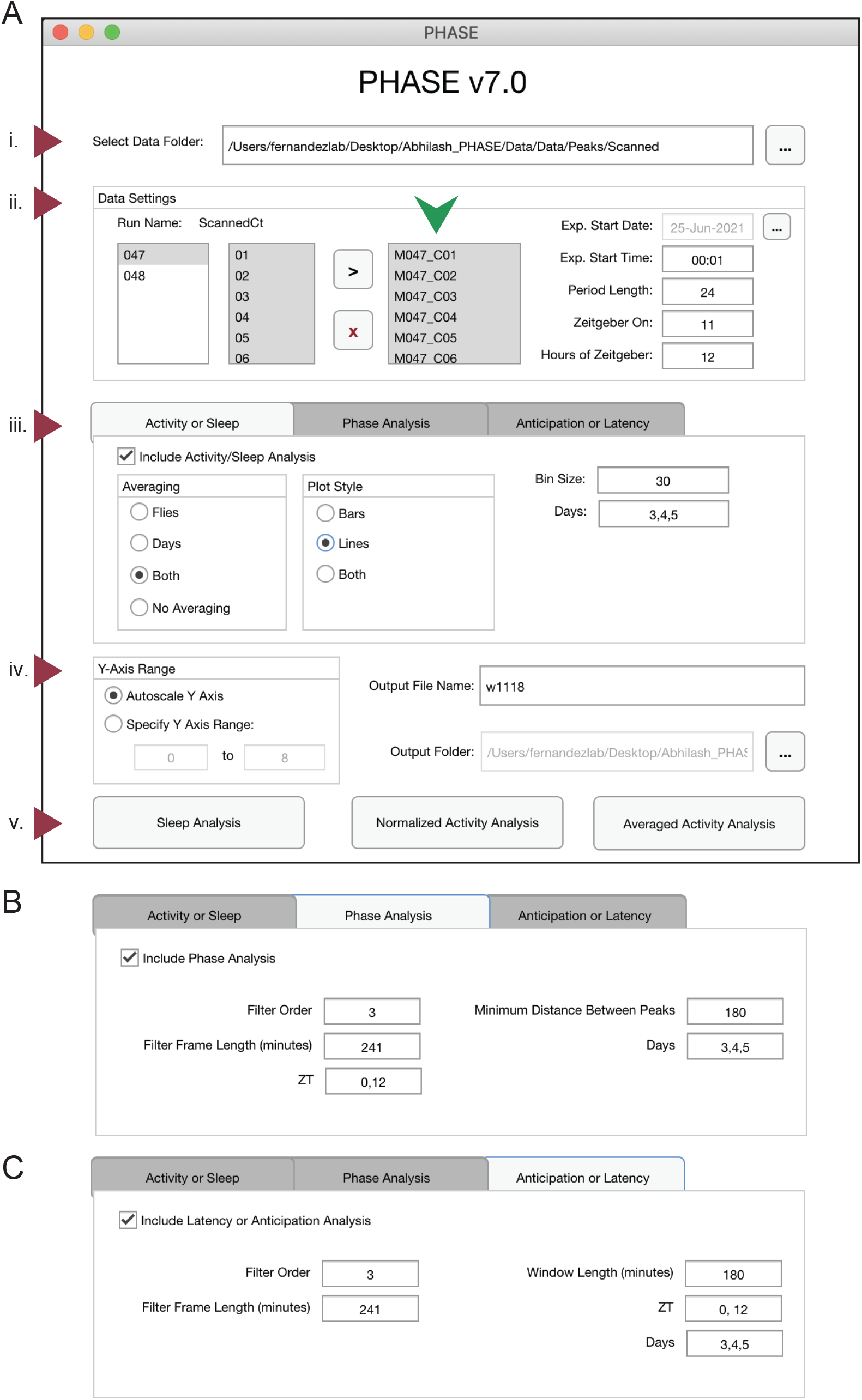
The PHASE graphical user interface. (A) The PHASE window as displayed on startup. The interface can be divided into five sections. (i) The “Select Data Folder” dialogue box: users click the button with three dots on the right to locate and open the folder containing one-minute scanned DAM channel files for analysis. PHASE automatically detects the monitor numbers and displays all available channels under “Run Name: ScannedCt” in the “Data Settings” section (ii). Users must move the channels to be analyzed by pushing them to the small display window to the right (green arrowhead) using the black “>“symbol. All channels within this box must then be selected for analysis to continue. Monitors must be analyzed one at a time. Experimental (Exp.) Start Date must be indicated by clicking the button with three dots in the top-right corner of section (ii). Exp. Start Time must be input in the HH:MM format using military time (00:00-23:59) and must start 1 minute after the intended experimental start time. In this example the experiment would begin at midnight on 25 June 2021. “Period Length” is the length of one full environmental cycle. This defaults to 24 in PHASE but can be adjusted for experiments using non-24-h environmental cycles. “Zeitgeber On” is the time at which experimental Zeitgeber (e.g., light, or high temperature) starts and is indicated as hours after Exp. Start Time. “Hours of Zeitgeber” indicates the duration of the zeitgeber ON time and is expressed in hours. In this example, the lights on period lasted 12-h per cycle, and lights turned on at 11:00 AM. (iii) The “Analysis” section has three tabs. Users can visualize and analyze activity/sleep profiles and export corresponding metrics using the “Activity or Sleep” tab; the timing of daily peaks of activity can be analyzed using the “Phase Analysis” tab (B); and the anticipation of environmental transitions can be analyzed using the “Anticipation or Latency” tab (C). These three analysis tabs can be run singly or in any combination. The “Phase Analysis” and “Anticipation or Latency” tabs have distinct interfaces and require the inputs shown in panels (B) and (C). To include these analyses in PHASE output files, the “Include…” box on the top-left of each tab must be checked (iii). For the “Activity or Sleep” analysis, users must also choose the way the data will be averaged (across days, across flies, or both), the plot style (lines or bars), the bin size (in minutes) and the days to be analyzed (separated by commas). In this example (A-iii), we have used 30-min bins and analyzed days 3, 4 and 5. The following inputs described are relevant for both the “Phase Analysis” and “Anticipation or Latency” tabs (B) and (C). The values of these inputs need not necessarily be the same for the two options. Filter Order and Filter Frame Length (minutes) inputs pertain to the Savitzky-Golay smoothing filter used for identifying peaks (see text for details). ZT (Zeitgeber Time) indicates the times of the day around which peaks must be identified (for “Phase Analysis”) or before which anticipation must be quantified (for “Anticipation or Latency” analysis), with ZT00 always defined as the onset of zeitgeber. Multiple ZT values can be provided, separated by commas, to characterize any number of consistently timed peaks or anticipation to environmental transitions within the time-series. Days should indicate the days over which phase and/or anticipation or latency must be estimated. These can be different from the days used as input in panel (A-iii). The “Phase Analysis” tab requires an additional input, “Minimum Distance Between Peaks” (B), which must be provided in minutes and is used by PHASE to identify multiple peaks outside this user-defined window. The “Anticipation or Latency” tab has an additional input, Window Length, which is expressed in minutes and sets the window of time prior to the chosen ZTs within which PHASE will estimate anticipation. The “Output” section (iv) has three dialogue boxes. The first, “Y-Axis Range” determines if the range of *y*-Axis values in the sleep/activity plots will be set automatically or by user-specified values. The remaining dialogue boxes require the user to the set “Output File Name,” and the “Output Folder,” which determines where output files will be saved. The three-dot button on the right of the Output Folder dialogue box is where the user navigates to the target. (v) The analysis execution buttons allow to either analyze Sleep, Normalized Activity or Averaged Activity (see text for details). All the analysis chosen in panels (A-iii, B or C) will be executed only when one of these three buttons are clicked.

### Activity and Sleep Analysis

PHASE allows users to view activity and sleep profiles in ways that facilitate easy visual inspection of inter-individual and cycle-to-cycle variations in temporal patterns (Figs. 2 and 3). It also computes and exports all underlying data to spreadsheets that also contain commonly used metrics for assessment of sleep and activity, such as the number of beam crossings (i.e., activity) and the number and duration of sleep bouts in the day and night-phases (see manual). Distinct types of sleep and activity plots can be generated in PHASE using different approaches (see Fig. 1A-iii): averaging data across flies, days, and/or both (Figs. 2A-C and 3A-C). When averaging across “Days” is chosen, one plot for every individual fly (i.e., channel) is generated wherein activity/sleep is averaged across all cycles indicated by the user (Figs. 2A and 3A). PHASE generates one PDF file for every 12 channels. Thus, the 32 channels of a standard DAM monitor are saved as three PDF files. When averaging across “Flies” is chosen, a single averaged plot of the activity or sleep timeseries is generated for the days indicated in 1A-iii and data are averaged across all the channels chosen by the user (Figs. 2B and 3B). When averaging across “Both,” a single plot of activity/sleep profile is generated, wherein data are averaged across all the days and flies indicated by the user (Figs. 2C and 3C).

**Figure 2:**
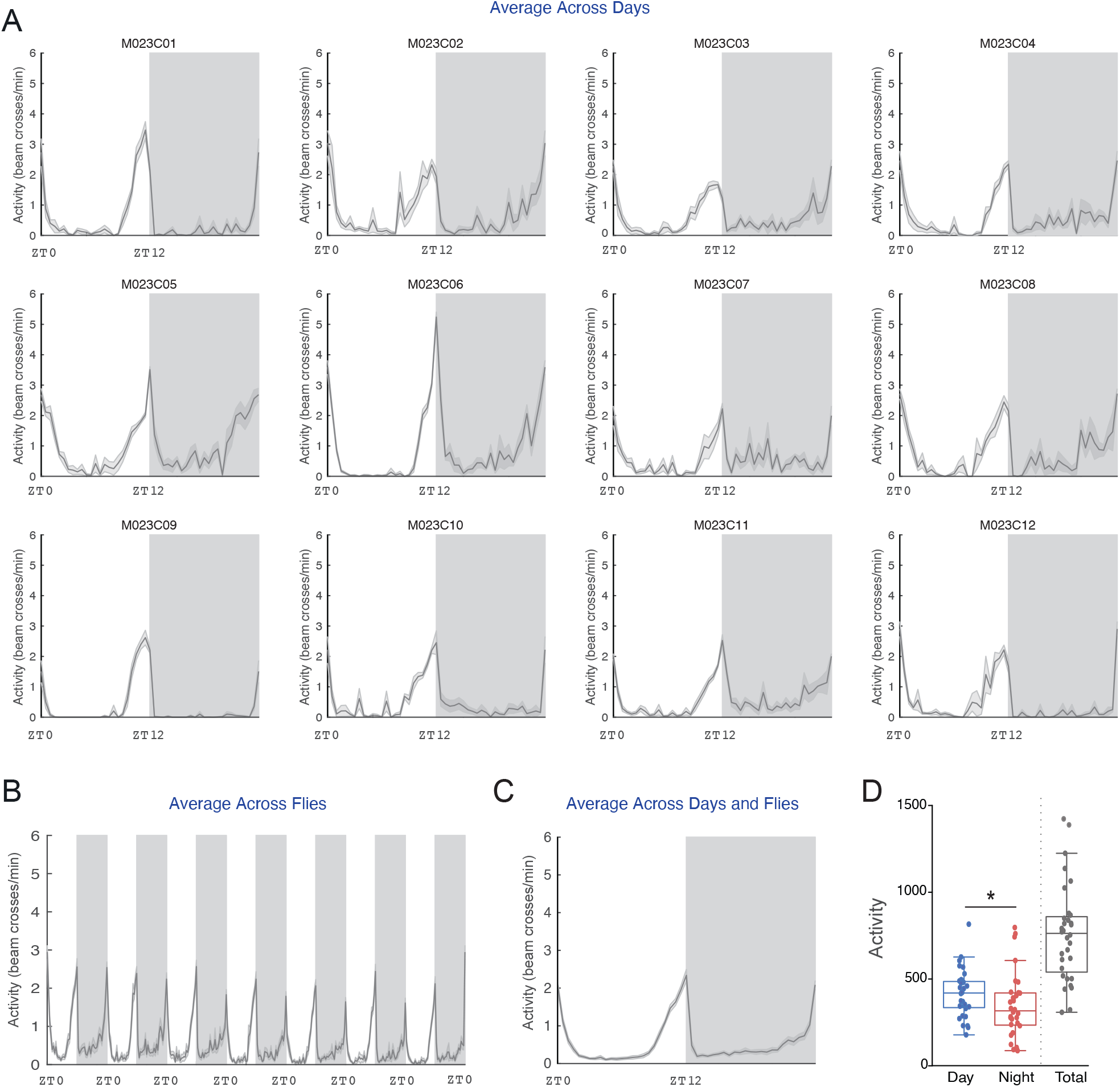
Averaged Activity Analysis” outputs from PHASE summarizing activity across flies and environmental cycles. Here, example activity time-courses are displayed for male *Canton-S* (*CS*) flies recorded under LD12:12 cycles at a constant 25 °C. (A) Averaged activity output from PHASE when time-series are averaged across days for 12 individual flies (see panel A-iii in Fig. 1). (B) Output from PHASE when time-series are averaged across all 32 flies (see panel A-iii in Fig. 1). (C) Output from PHASE when time-series are averaged across both days and flies (see panel A-iii in Fig. 1). Gray shaded regions in panels (A) through (C) indicate the scotophase (i.e., darkness) of the light/dark cycle. When these analyses are executed, corresponding spreadsheets are generated (see user manual for details), which can be used to visualize and perform statistical tests on activity parameters. One such example is shown in (D), wherein total day-time and night-time activity in *CS* flies is compared. As described previously, day-time activity is significantly higher than night-time activity as judged by a Wilcoxon’s matched pairs test implemented in R (*V* = 395; *p* = 0.013; also indicated on the plot using an asterisk). Also shown, for relative comparison, is the total activity counts across one full cycle (not included in any statistical comparisons). Note that such plots can be visualized as normalized time-series using the “Normalized Activity Analysis” output button (Fig. 1A-v).

**Figure 3:**
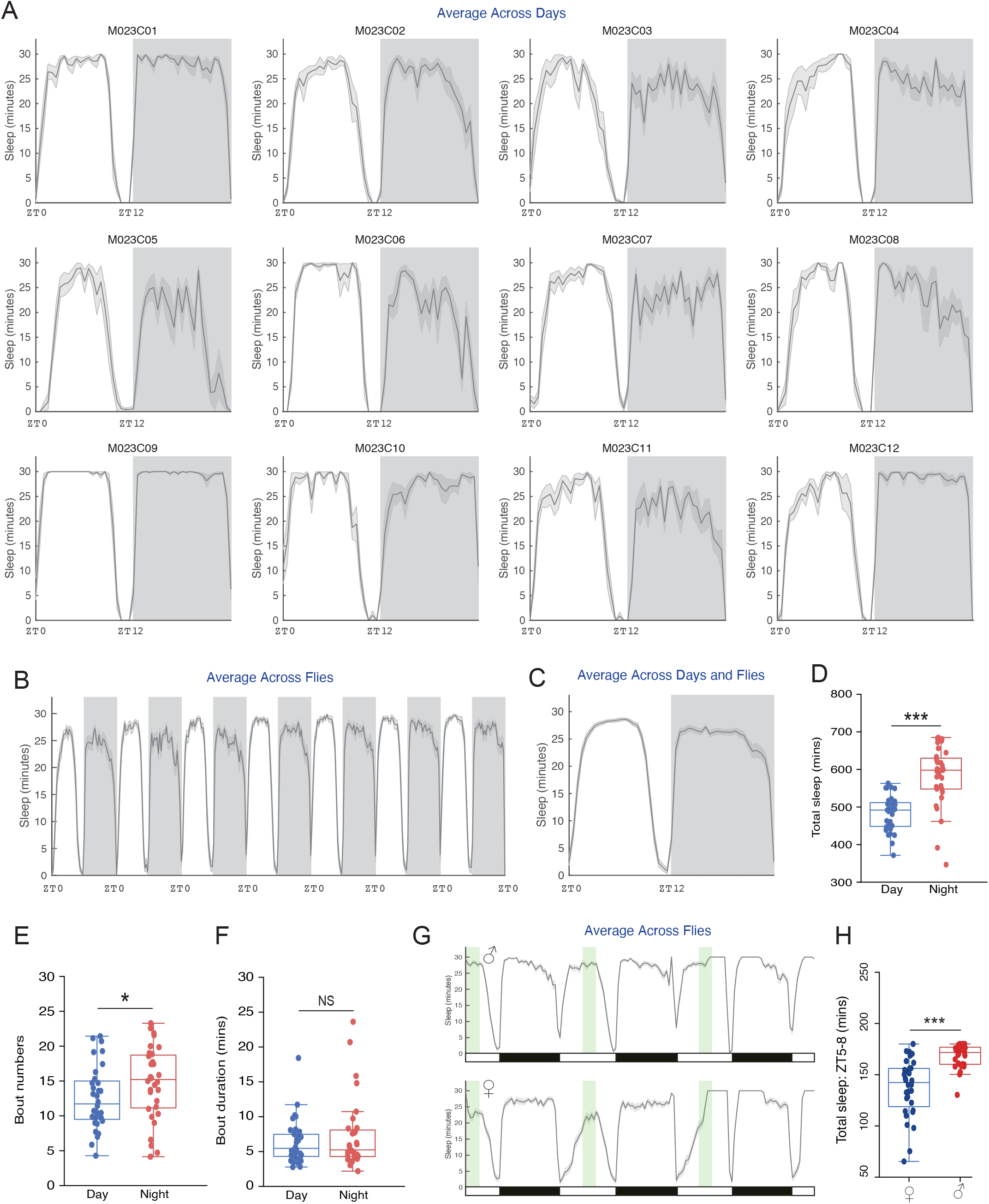
Sleep Analysis” outputs from PHASE. Shown here are plots of sleep, measured as minutes/user-defined bin, averaged across days for 12 individual flies (A), across all 32 flies (B) and across both days and flies (C) in the same manner as that shown for averaged activity, using the same time-series used for the analysis described in Fig. 2 (see Figs. 1A-iii and 2). Gray shaded regions in panels (A) through (C) indicate the scotophase of the light/dark cycle. The exported spreadsheets from the PHASE’s sleep analysis were used to compare total sleep, sleep bout numbers and duration across day and night in male *CS* flies. (D) As previously described, total minutes of sleep is higher during the night (*V* = 15; *p* = 6.38e-08). (E) The number of sleep bouts was significantly higher during the night (*V* = 153.5; *p* = 0.04). (F) Sleep bout duration was not significantly different between day and night (*V* = 241; *p* = 0.68). All statistical inferences in panels (D), (E) and (F) were drawn based on Wilcoxon’s matched pairs tests implemented in R. (G) Also shown are plots of sleep, measured as minutes/user-defined bin, averaged across flies for males (top) and females (bottom) for three consecutive LD cycles. To select a three-hour window around ZT06, three-hours was defined as the “Hours of Zeitgeber” and “Zeitgeber On” time was changed accordingly in the “Data Settings” input panel (see Fig. 1A-ii). This modified daytime window is shown in green, while the actual LD cycle during the experiment is indicated by the black and white bars underneath the plot. In this window we quantified total minutes of sleep for females and males. (H) As expected from previous work, total minutes of sleep in this three-hour window was significantly higher in males than in females as judged by a Mann-Whitney U test implemented in R (*W* = 133.5; *p* = 3.84e-07). For all panels: * < 0.05, ** < 0.01, *** < 0.001, and NS (Not Significant).

As previously described by many investigators, activity was significantly higher during the day in wild-type *Canton S* (*CS*) male flies under LD12:12 (Fig. 2D) and total sleep was higher during the night (Hamblen-Coyle et al., 1992; Majercak et al., 1999; Hendricks et al., 2000; Shaw et al., 2000). Increased nighttime sleep was accompanied by increased bout numbers at night (Fig. 3E) with no differences in the bout duration between night and day (Fig. 3F). Sexual dimorphism in sleep is well documented (Huber et al., 2004; Andretic and Shaw, 2005), with females sleeping less during the day than males. We quantified sleep within a three-hour window around mid-day (ZT05-ZT08) to highlight how window-specific analysis of sleep/activity can be carried out using PHASE (Fig. 3G). As expected, this analysis revealed that daytime sleep was significantly higher in males than in females (Fig. 3H).

### Savitzky-Golay filtering for the quantification of behavioral phase

One of the earliest methods used to objectively characterize the timing of activity peaks in *Drosophila* was described by (Hamblen-Coyle et al., 1992). These authors employed a Butterworth filter to remove high-frequency components of beam crossing time-course data and used a search algorithm to look for peaks within specific windows of interest from the filtered data. Other objective measures have included the use of a non-recursive digital filter (Helfrich-Förster, 1998), and dual smoothing (non-recursive digital filter + moving averages) (Rieger et al., 2003). Recent studies have used various alternative methods of peak detection (Potdar and Vasu, 2012; De et al., 2013; Das et al., 2015; Liang et al., 2019), which supports the need for a robust peak detection method and software to implement it. While it is useful to use smoothed data to quantify the timing of behavioral peaks, there are limitations to using filters like the Butterworth because such filters work well only for stationary time-series (wherein frequency and amplitude are maintained from cycle-to-cycle). Behavioral rhythms are typically non-stationary and therefore require tools that work well for such time-series (Leise, 2013; Leise, 2017). One such digital filter is the Savitzky-Golay (SG) filter (Savitzky and Golay, 1964; Schafer, 2011). The SG filter is known for its ability to preserve amplitude of the signal while de-noising the data set, thereby making it extremely useful for the identification and characterization of peaks and the estimation of their phases. While it has been used in other sub-disciplines of biology (Azami et al., 2012; Chu et al., 2014), it has not, to our knowledge, been applied to circadian data.

Mathematically, the SG filter fits a polynomial of order *p*, to a window of size *n* over the original timeseries using linear least-squares. This window of the timeseries then slides across the entire length of the timeseries, fitting a polynomial of order *p* for each window. In the case of equally spaced data, as is the case with *Drosophila* activity and sleep time-series, there exists an analytical solution to this set of least-squares equations as a single set of convolution coefficients, thereby making this a fast and computationally straightforward filter. These coefficients can then be used to derive signal estimates (e.g., activity or sleep) at a given time *t*. This smoothed, estimated signal can then be used to make peak calls and characterize anticipation of environmental transitions (see below). For its implementation in PHASE, it is important to note that the sliding window must consist of an odd number of timepoints, whose units must be in minutes.

To demonstrate the effect of polynomial order and window length on smoothing, we used SG filters with varying values of *p* and *n* and overlaid smoothed estimates over the raw data (Suppl. Fig. 1). These results show that, for a given filter order, increasing window lengths result in smoother time-series. Furthermore, for a given window length, increasing the filter order increases the ability of the filter to capture nuances of the time-series (Suppl. Fig. 1). Therefore, the most useful filter order and frame-length will depend on the nature of both the raw data and the question being posed, which can be established using an iterative process to determine the most appropriate parameters.

We used a filter order of three and a frame-length of 241 minutes to estimate peak times in the analysis presented here to illustrate how this filter can be used to characterize activity peaks in *Drosophila* behavioral time-series. Filtering with these parameters captures the activity patterns of wild-type *CS* males entrained to three LD cycles of differing daylengths well (Fig. 4A-left). For equinox and short-day cycles (LD 8:16, 8 hours of light and 16 hours of darkness), the morning peaks of activity immediately follow lights-on but occur about an hour after lights-on under long-day conditions (Fig. 4B). Evening peaks of activity are clearly adjusted to occur just before dusk (Fig. 4C) in *CS* males, as previously described (e.g., Qiu and Hardin, 1996). In contrast, *per*^*01*^ mutants, which lack a functional molecular clock, are characterized by evening peaks that occur after lights-off under all day-lengths (Figs. 4A-right and 4C), suggesting that these are masked peaks of activity and not clock-driven activity peaks. This illustrates the utility of the SG filter for determining if behavioral peaks anticipate or follow environmental transitions.

**Figure 4:**
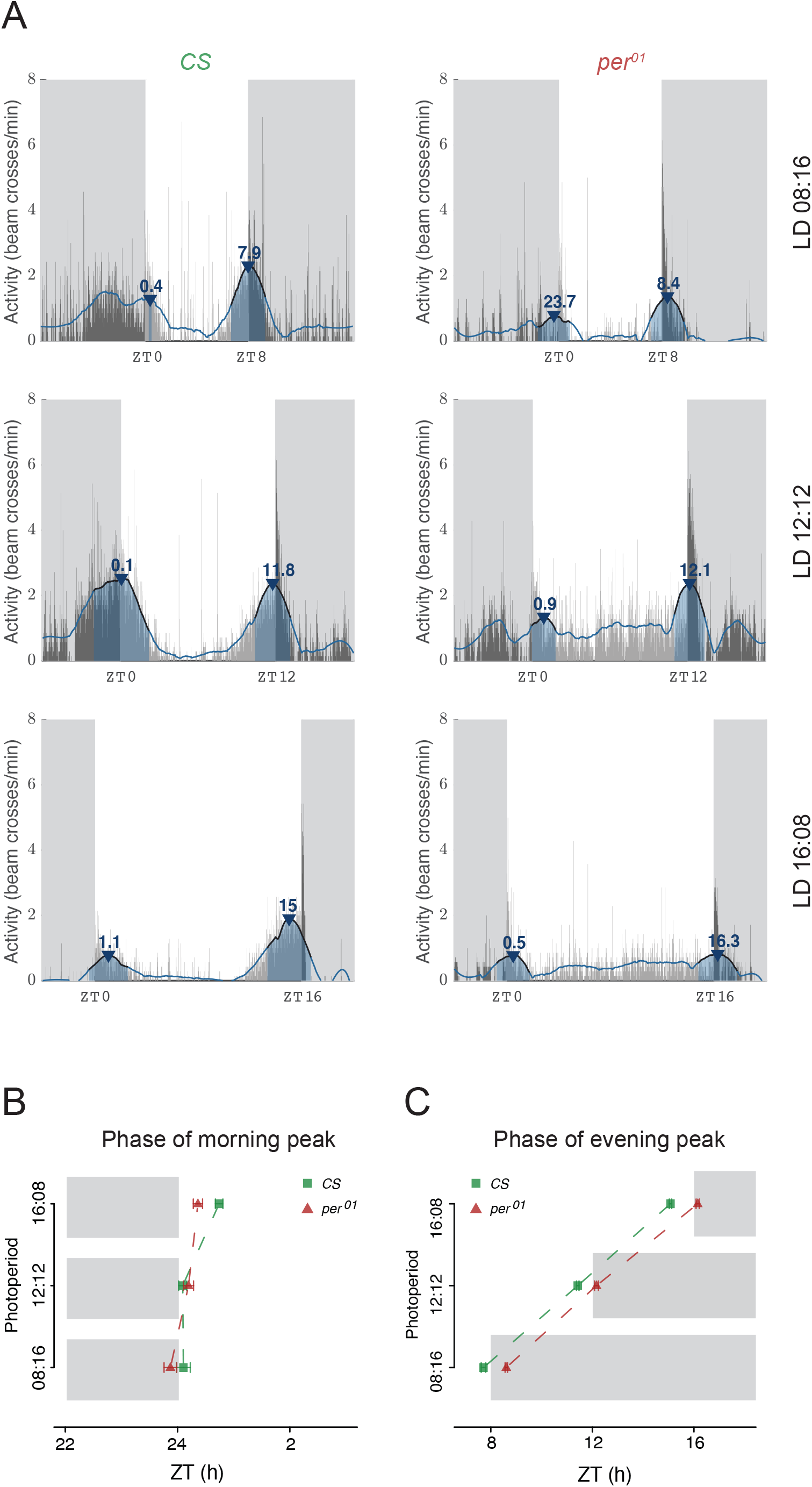
Phase Analysis” outputs from PHASE. (A) Representative activity of *Canton-S* (*CS*) male flies under LD08:16 (top-left), LD12:12 (middle-left), and LD16:08 (bottom-left), and *per*^*01*^ flies in the same three regimes (right). In all cases, the thin gray vertical bars show the activity of the fly in one-minute bins averaged across all days chosen by the user. The blue line is the smoothed waveform produced by the Savitzky-Golay filter (see text for details). Peaks are identified based on the smoothed waveform represented by the blue line and are indicated by the blue arrowheads. Annotated above the arrowheads are the phases of the identified peaks expressed as ZTs. The blue shaded region under the arrowhead is the width of the peak. Also shown are mean±SEM (Standard Errors of the Mean) of the phases of morning (B) and evening (C) peaks of activity for *CS* and *per*^*01*^ flies under the three photoperiods. Gray shaded regions in both (B) and (C) indicate the scotophase of the light/dark cycle in their respective photoperiod regimes.

We used a second fly mutant to demonstrate the utility of the SG filter in the characterization of behavioral timing. As previously shown, *pdf*^*01*^ mutants have an advanced evening peak of activity and a reduced or absent morning peak of activity under LD cycles (Renn et al., 1999). Comparing the top and bottom rows of Fig. 5A reveals that the morning peaks of activity are highly reduced in *pdf*^*01*^ flies compared to their genetic control background, *w*^*1118*^. We extracted peak information with PHASE using the order and window length values described above and quantified both the phases and heights of activity peaks for both genotypes. As expected from previous work (Renn et al., 1999), the morning peaks of *pdf*^*01*^ mutants were delayed (Fig. 5B) and their heights significantly reduced (Fig. 5C), whereas the evening peak phases were significantly advanced (Fig. 5D) and their heights significantly reduced compared to *w*^*1118*^ controls (Fig. 5E). These examples further illustrate the utility of smoothing time-series data with the SG filter to characterize the timing and magnitude of activity peaks. A similar approach can be used to estimate and characterize peaks of sleep in flies using PHASE.

**Figure 5:**
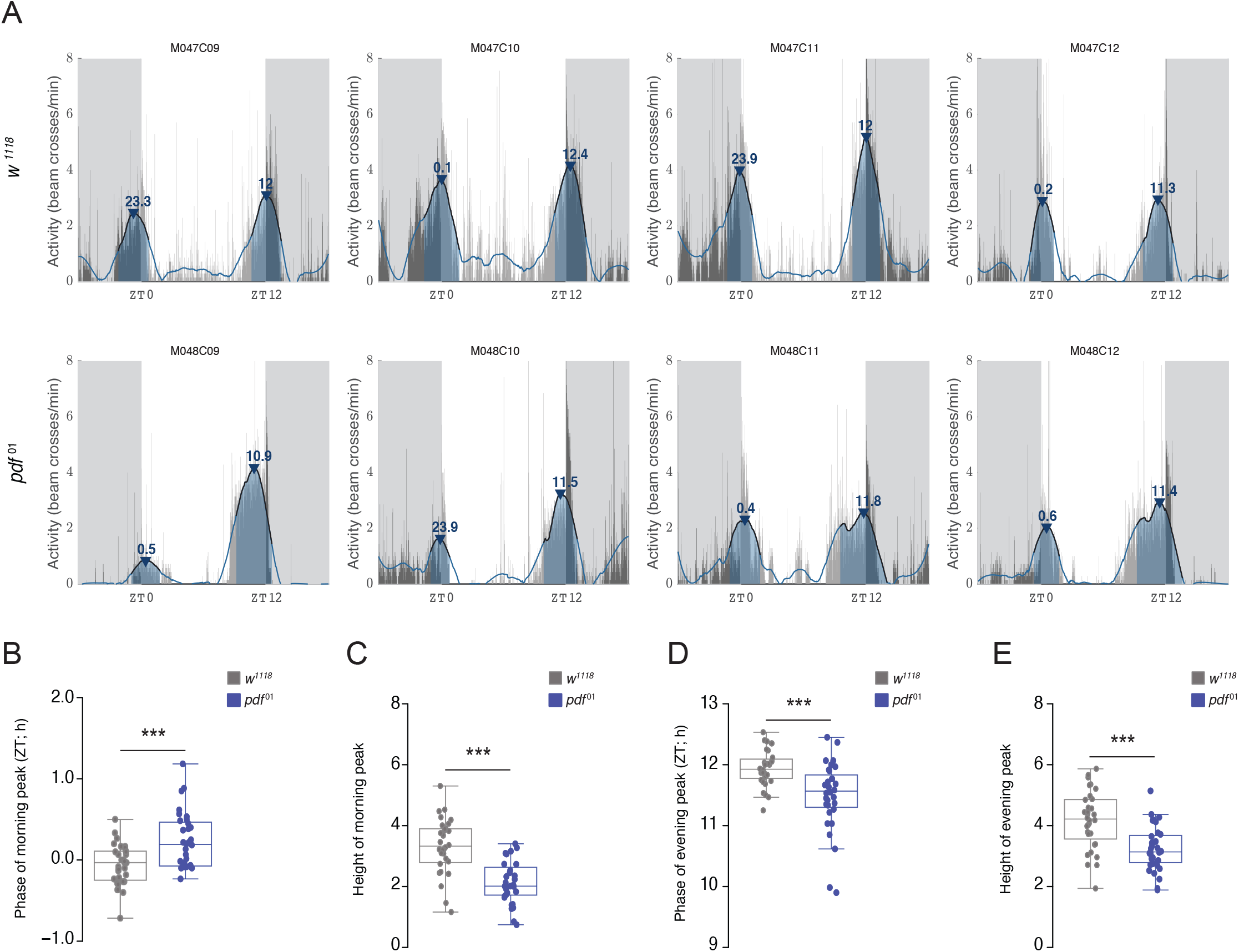
Characterization of activity peaks in *pdf*^*01*^ mutants. (A) Representative activity profiles of four males each of *w*^*1118*^ controls (top) and *pdf*^*01*^ mutants (bottom) overlaid with SG-filtered waveforms show advanced evening peaks and reduced morning peaks in *pdf*^*01*^ flies. We quantified phases and heights of both morning and evening activity peaks and found that phase of morning peak is significantly delayed in *pdf*^*01*^ flies (B; *W* = 264.5; *p* = 0.0009), and the height of morning peak is significantly lower compared to *w*^*1118*^ flies (C; *W* = 879; *p* = 1.581e-07). On the other hand, the phase of evening peak is significantly advanced (D; *W* = 794.5; *p* = 0.0001521) and its height significantly lower (E; *W* = 800; *p* = 6.718e-05) in *pdf*^*01*^ mutants compared to *w*^*1118*^ flies. Comparisons were done using Mann-Whitney U tests implemented on R. * < 0.05, ** < 0.01, *** < 0.001, and NS (Not Significant).

### The Use of Slopes for Estimating Anticipation

The behavioral anticipation of predictable environmental transitions depends on a functional and entrained circadian system. One of the first methods of estimating anticipation was proposed by (Stoleru et al., 2004) who used an estimate of anticipation that was directly proportional to the gradual, stepwise increase in activity prior to environmental transitions, inversely proportional to the startle response due to the transition. Only a few studies have made use of this method (Stoleru et al., 2004; Stoleru et al., 2007; Sheeba et al., 2010). A simpler approach to the estimation of anticipation was proposed by (Harrisingh et al., 2007), who used the ratio of activity in a three-hour window before a transition of interest and the activity in a six-hour window before the same transition. The expected value of this metric in case of a complete lack of anticipation is 0.5. As anticipation increases, the value of this metric is also expected to increase. This approach has been used in many studies (Im and Taghert, 2010; Sheeba et al., 2010; Potdar and Vasu, 2012; De et al., 2013; Prakash et al., 2017; Baik et al., 2018; Schlichting et al., 2019). A recently published approach to quantifying anticipation made use of slopes from least-squares linear regression modeling of the increase in activity before specific environmental transitions (Fernandez et al., 2020). While both the above discussed measures of anticipation can be easily calculated from the activity data exported by PHASE, the slope method requires additional steps. We therefore implemented slope calculations into our analysis package to ease the use of this approach for *Drosophila* behavioral time-series data.

For the calculation of slopes, PHASE allows users to choose the duration of the time-window and the temporal resolution of time-series data sets to conduct linear regression analysis. Using this method, we compared the activity of wild-type (*CS*) and *per*^*01*^ mutant males in the hours before lights-on and light-off under LD12:12 cycles. Best fitting lines for each fly reveal that *CS* flies (Fig. 6A) display steeper increases in activity (i.e., higher slopes) before both the morning and evening transitions compared to *per*^*01*^ mutants (Fig. 6B). We quantified morning and evening slopes for every fly and compared average slope values for each genotype. The results revealed that, as expected, *per*^*01*^ mutants do not display significantly positive slopes in the hours leading to light transitions, indicating that there is no morning or evening anticipation, consistent with the absence of circadian timekeeping (Fig. 6C and legend therein). Furthermore, *CS* flies consistently displayed significantly positive slopes that were on average much higher than those of *per*^*01*^ flies, consistent with the presence of robust anticipatory activity in *CS* controls (Fig.6C). It is important to note here that this functionality is available only for the “Averaged Activity Analysis” option (see Fig. 1A-v) and not for the “Normalized Activity Analysis” option.

**Figure 6:**
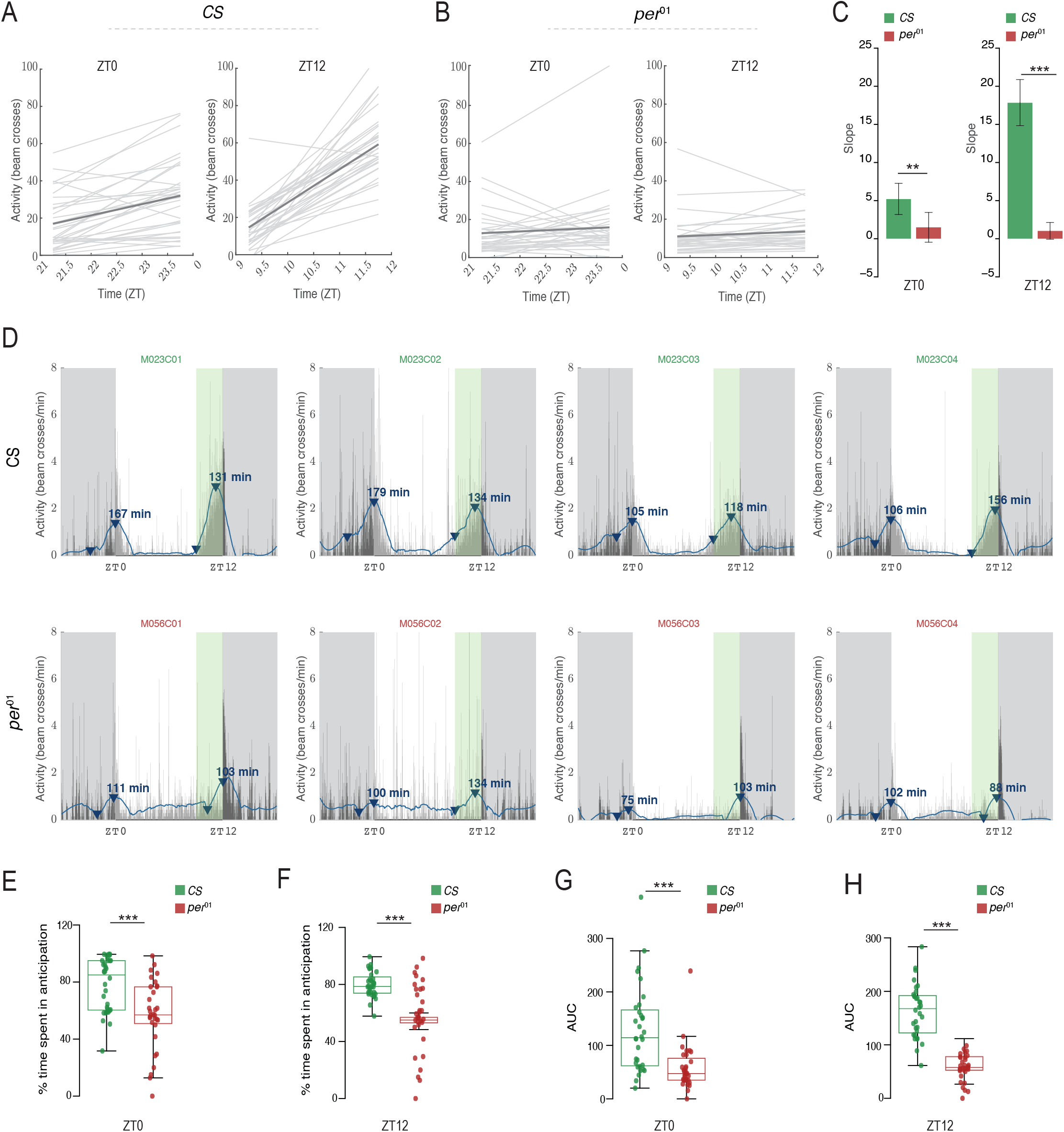
Anticipation or Latency” outputs from PHASE. Plots indicating slopes of activity levels in user-defined windows for individual male *Canton-S* (*CS)* (A) and *per*^*01*^ (B). Light gray lines represent least squares linear regressions for individual flies, and the dark gray line represents the averaged regression across all flies. (C) The slope values represented in panels (A) and (B) are exported within a PHASE-generated spreadsheet and are used to making comparisons between genotypes or treatments. A population with no anticipation would be expected to show a slope of zero. We performed single-sample *t*-tests to determine if the mean slope for each genotype was different from zero. *CS* flies have slope values that are significantly greater than zero in both the morning (left; *t*_*31*_ = 5.2057; *p* = 1.191e-05) and evening (right; *t*_*31*_ = 12.051; *p* = 3.118e-13) windows. On the other hand, *per*^*01*^ flies did not have significant anticipation in the morning (left; *t*_*31*_ = 1.564; *p* = 0.128) or evening (right; *t*_*31*_ = 1.928; *p* = 0.063) window. The error bars on these plots are 95% CI based on the reported *t*-tests. Therefore, any mean with error bars that cross the zero-mark is not different from zero. We also performed Welch’s *t*-tests to compare anticipation values between *CS* and *per*^*01*^ flies in both windows. We found that anticipation in the morning (left; *t*_*61*.*87*_ = 2.686; *p* = 0.009) and evening (right; *t*_*39*.*04*_ = 10.669; *p* = 3.912e-13) windows is significantly greater in *CS* flies as compared to *per*^*01*^ mutants. (D) Representative activity profiles (channels 1 through 4) of *Canton-S* (CS; top) and *per*^*01*^ (bottom) flies under LD12:12. The thin vertical bars represent the activity of the fly in 1-min bins averaged over all days chosen by the user. The blue line is the smoothed waveform after applying a Savitzky-Golay filter (see text for details). Dark gray shaded region is the scotophase of the light/dark cycle. The green shaded region in the plot (clearly visible before ZT12) is the user-defined window in which anticipation/latency is computed. The blue arrowheads show the minimum and maximum value of activity on the smoothed waveform in the user-defined window. Note that the arrowhead is not displayed for smoothed data points that are negative. The arrowhead marking the maximum value is annotated with the duration of anticipation or latency (in minutes; in other words, duration between the minimum and maximum value). (E-F) Percentage of time spent in anticipation is significantly higher in *CS* than *per*^*01*^ flies in both the morning (E; _*1*_*χ*^*2*^ = 13.052; *p* = 0.0003) and evening (F; _*1*_*χ*^*2*^ = 40.713; *p* = 1.763e-10) windows. Similarly, areas under the curve (AUC) were also significantly higher in *CS* than *per*^*01*^ flies in morning (G; _*1*_*χ*^*2*^ = 19.043; *p* = 1.278e-5) and evening (H; _*1*_*χ*^*2*^ = 37.324; *p* = 1e-09) windows. All inferences in panels (B) through (E) were made using Kruskal-Wallis tests implemented in R. For all panels: * < 0.05, ** < 0.01, *** < 0.001, and NS (Not Significant). Note that a similar approach can be used on sleep data to summarize increases in sleep following environmental transitions as a means of quantifying the latency of sleep (see manual).

The SG filter can also identify the highest and lowest points of activity in the smoothed data within the user-defined window (Fig. 6D). PHASE uses these values to estimate the duration of anticipatory activity and the area under the curve between the minimum and maximum data points identified by PHASE (Fig. 6D). Based on these measurements (see manual for details), we found that the percent time spent in anticipation was significantly higher in *CS* controls compared to *per*^*01*^ mutants for both morning (Fig. 6E) and evening (Fig. 6F) transitions. We also found that the area under the curve was significantly larger for *CS* controls for both transitions (Figs. 6G and H), consistent with significantly reduced anticipation in *per*^*01*^ mutants. Additionally, the slope method is utilized by PHASE to calculate anticipation of environmental transitions for “Normalized Activity” and latency of sleep following environmental transitions using the smoothed normalized activity and sleep data, respectively (see manual for details). We note that each method of measuring anticipation has its own set of limitations and the ability to implement multiple measures in a program like PHASE will be helpful for researchers interested in quantitively characterizing anticipation in fly behavioral time-series.

In conclusion, we offer PHASE as a platform to support the visualization and analysis of behavioral time-series through user-defined, window-based measurements of activity and sleep data. Furthermore, PHASE implements a powerful smoothing filter to model fly activity and sleep rhythms, which allows for the quantitative characterization of activity peaks and anticipation, a feature that is not currently widely available. All codes for PHASE are open-source and available at: https://github.com/ajlopatkin/PHASE. The software bundle for installation can be downloaded from the same link. A detailed user manual is available as supplementary online material. This user manual contains detailed information regarding the interface, suggestions to make analyses easier, details about the graphs generated, and descriptions of common errors and how to troubleshoot them, all of which we hope will make this program useful for the circadian fly community and support future breakthroughs in our understanding of circadian entrainment.

## Supporting information

Supplemental Figure and Manual

## Acknowledgments

This work was supported by National Institutes of Health (NIH) NINDS grant (R01NS077933) and an NSF IOS grant (1354046) to OTS, an NIH NINDS grant (R01NS118012) to MPF and OTS, and start-up funds from Barnard College to MPF. We thank members of the Fernandez and Shafer labs for feedback on the PHASE software.

## References

Abhilash L and Sheeba V (2019) RhythmicAlly: Your R and Shiny–Based Open-Source Ally for the Analysis of Biological Rhythms. J Biol Rhythms 074873041986247. doi: 10.1177/0748730419862474.

Andretic R and Shaw P (2005) Essentials of sleep recordings in Drosophila: moving beyond sleep time. Meth Enzymol 393:759–772. doi: 10.1016/S0076-6879(05)93040-1.

Azami H, Mohammadi K and Bozorgtabar B (2012) An Improved Signal Segmentation Using Moving Average and Savitzky-Golay Filter. 2012:39–44. doi: 10.4236/JSIP.2012.31006.

Baik L, Recinos Y, Chevez J and Holmes T (2018) Circadian modulation of light-evoked avoidance/attraction behavior in Drosophila. PLoS One. doi: 10.1371/JOURNAL.PONE.0201927

Cenek L, Klindziuk L, Lopez C, McCartney E, Martin Burgos B, Tir S, Harrington M and Leise T (2020) CIRCADA: Shiny Apps for Exploration of Experimental and Synthetic Circadian Time Series with an Educational Emphasis. J Biol Rhythms 35:214–222. doi: 10.1177/0748730419900866.

Chiu JC, Low KH, Pike DH, Yildirim E and Edery I (2010) Assaying locomotor activity to study circadian rhythms and sleep parameters in Drosophila. J of Vis Exp: JoVE. doi: 10.3791/2157

Chu L, Liu GH, Huang C and Liu QS (2014) Phenology detection of winter wheat in the Yellow River delta using MODIS NDVI time-series data. 2014 The 3rd International Conference on Agro-Geoinformatics, Agro-Geoinformatics 2014. doi: 10.1109/AGRO-GEOINFORMATICS.2014.6910664

Cichewicz K and Hirsh J (2018) ShinyR-DAM: a program analyzing Drosophila activity, sleep and circadian rhythms. Comm Biol 2018 1:1 1:1–5. doi: 10.1038/s42003-018-0031-9.

Das A, Holmes TC and Sheeba V (2015) dTRPA1 Modulates Afternoon Peak of Activity of Fruit Flies Drosophila melanogaster. PLoS One 10:e0134213. doi: 10.1371/JOURNAL.PONE.0134213.

De J, Varma V, Saha S, Sheeba V and Sharma VK (2013) Significance of activity peaks in fruit flies, Drosophila melanogaster, under seminatural conditions. Proc Natl Acad Sci USA 110:8984–9. doi: 10.1073/pnas.1220960110.

Fernandez MP, Pettibone HL, Bogart JT, Roell CJ, Davey CE, Pranevicius A, Huynh K v., Lennox SM, Kostadinov BS and Shafer OT (2020) Sites of Circadian Clock Neuron Plasticity Mediate Sensory Integration and Entrainment. Curr Biol 30:2225–2237.e5. doi: 10.1016/J.CUB.2020.04.025.

Geissmann Q, Rodriguez LG, Beckwith EJ and Gilestro GF (2019) Rethomics: An R framework to analyse high-throughput behavioural data. PLoS One 14:e0209331. doi: 10.1371/JOURNAL.PONE.0209331.

Grima B, Chélot E, Xia R and Rouyer F (2004) Morning and evening peaks of activity rely on different clock neurons of the Drosophila brain. Nature 431:869–873. doi: 10.1038/nature02935.

Hamblen-Coyle MJ, Wheeler DA, Rutila JE, Rosbash M and Hall JC (1992) Behavior of period-altered circadian rhythm mutants of Drosophila in light:dark cycles (Diptera: Drosophilidae). J Insect Behav 5:417–446. doi: 10.1007/BF01058189.

Hammad G, Reyt M, Beliy N, Baillet M, Deantoni M, Lesoinne A, Muto V and Schmidt C (2021) pyActigraphy: Open-source python package for actigraphy data visualization and analysis. PLoS Comput Biol 17:e1009514. doi: 10.1371/JOURNAL.PCBI.1009514.

Hardin PE (2011) Molecular genetic analysis of circadian timekeeping in Drosophila. Adv Genet 141–173.

Harrisingh MC, Wu Y, Lnenicka GA and Nitabach MN (2007) Intracellular Ca2+ Regulates Free-Running Circadian Clock Oscillation In Vivo. J Neurosci 27:12489–12499. doi: 10.1523/JNEUROSCI.3680-07.2007.

Helfrich-Förster C (1998) Robust circadian rhythmicity of Drosophila melanogaster requires the presence of lateral neurons: a brain-behavioral study of disconnected mutants. J Comp Physiol A 182:435–453. doi: 10.1007/S003590050192.

Helfrich-Förster C (2020) Light input pathways to the circadian clock of insects with an emphasis on the fruit fly Drosophila melanogaster. J Comp Physiol A 206:259. doi: 10.1007/S00359-019-01379-5.

Hendricks J, Finn S, Panckeri K, Chavkin J, Williams J, Sehgal A and Pack A (2000) Rest in Drosophila is a sleep-like state. Neuron 25:129–138. doi: 10.1016/S0896-6273(00)80877-6.

Huber R, Hill S, Holladay C, Biesiadecki M, Tononi G and Cirelli C (2004) Sleep homeostasis in Drosophila melanogaster. Sleep 27:628–639. doi: 10.1093/SLEEP/27.4.628.

Im S and Taghert P (2010) PDF receptor expression reveals direct interactions between circadian oscillators in Drosophila. J Comp Neurol 518:1925–1945. doi: 10.1002/CNE.22311.

Kostadinov B, Lee Pettibone H, Bell EV, Zhou X, Pranevicius A, Shafer OT and Fernandez MP (2021) Open-source computational framework for studying Drosophila behavioral phase. STAR Protocols 2:100285. doi: 10.1016/J.XPRO.2020.100285.

Leise TL (2017) Analysis of Nonstationary Time Series for Biological Rhythms Research: http://dx.doi.org/101177/0748730417709105 32:187–194. doi: 10.1177/0748730417709105.

Leise TL (2013) Wavelet analysis of circadian and ultradian behavioral rhythms. J Circ Rhythms 11:5. doi: 10.1186/1740-3391-11-5.

Liang X, Ho MCW, Zhang Y, Li Y, Wu MN, Holy TE and Taghert PH (2019) Morning and Evening Circadian Pacemakers Independently Drive Premotor Centers via a Specific Dopamine Relay. Neuron 102:843–857.e4. doi: 10.1016/J.NEURON.2019.03.028.

Majercak J, Sidote D, Hardin PE and Edery I (1999) How a circadian clock adapts to seasonal decreases in temperature and day length. Neuron 24:219–230.

Moore-Ede MC, Sulzman FM and Fuller CA (Charles A (1982) The clocks that time us : physiology of the circadian timing system. Harvard University Press

Pittendrigh CS (1981) Circadian Systems: Entrainment. Biological Rhythms. Springer US, Boston, MA, pp 95–124

Potdar S and Vasu S (2012) Large Ventral Lateral Neurons Determine the Phase of Evening Activity Peak across Photoperiods in Drosophila melanogaster. J Biol Rhythms 27:267–279. doi: 10.1177/0748730412449820.

Prakash P, Nambiar A and Sheeba V (2017) Oscillating PDF in termini of circadian pacemaker neurons and synchronous molecular clocks in downstream neurons are not sufficient for sustenance of activity rhythms in constant darkness. PLoS One 12:e0175073. doi: 10.1371/journal.pone.0175073.

Qiu J and Hardin PE (1996) per mRNA cycling is locked to lights-off under photoperiodic conditions that support circadian feedback loop function. Mol Cell Biol 16:4182–4188. doi: 10.1128/MCB.16.8.4182.

Renn SCP, Park JH, Rosbash M, Hall JC and Taghert PH (1999) A pdf neuropeptide gene mutation and ablation of PDF neurons each cause severe abnormalities of behavioral circadian rhythms in Drosophila. Cell 99:791–802. doi: 10.1016/S0092-8674(00)81676-1.

Rieger D, Stanewsky R and Helfrich-Förster C (2003) Cryptochrome, compound eyes, Hofbauer-Buchner eyelets, and ocelli play different roles in the entrainment and masking pathway of the locomotor activity rhythm in the fruit fly Drosophila melanogaster. J Biol Rhythms 18:377–391. doi: 10.1177/0748730403256997.

Roenneberg T, Dragovic Z and Merrow M (2005) Demasking biological oscillators: properties and principles of entrainment exemplified by the Neurospora circadian clock. Proc Natl Acad Sci USA 102:7742–7. doi: 10.1073/pnas.0501884102.

Savitzky Abraham and Golay MJE (1964) Smoothing and Differentiation of Data by Simplified Least Squares Procedures. Anal Chem 36:1627–1639. doi: 10.1021/AC60214A047.

Schafer RW (2011) What is a savitzky-golay filter? IEEE Signal Processing Magazine 28:111–117. doi: 10.1109/MSP.2011.941097.

Schlichting M, Díaz MM, Xin J and Rosbash M (2019) Neuron-specific knockouts indicate the importance of network communication to Drosophila rhythmicity. eLife. doi: 10.7554/eLife.48301

Schmid B, Helfrich-Förster C and Yoshii T (2011) A New ImageJ Plug-in “ActogramJ” for Chronobiological Analyses. J Biol Rhythms 26:464–467. doi: 10.1177/0748730411414264.

Shafer OT and Keene AC (2021) The Regulation of Drosophila Sleep. Curr Biol 31:R38–R49. doi: 10.1016/J.CUB.2020.10.082.

Shaw PJ, Cirelli C, Greenspan RJ and Tononi G (2000) Correlates of Sleep and Waking in Drosophila melanogaster. Science 287:1834–1837. doi: 10.1126/SCIENCE.287.5459.1834.

Sheeba V, Fogle KJ and Holmes TC (2010) Persistence of Morning Anticipation Behavior and High Amplitude Morning Startle Response Following Functional Loss of Small Ventral Lateral Neurons in Drosophila. PLoS One 5:e11628. doi: 10.1371/JOURNAL.PONE.0011628.

Stoleru D, Nawathean P, Fernández M de la P, Menet JS, Ceriani MF and Rosbash M (2007) The Drosophila Circadian Network Is a Seasonal Timer. Cell 129:207–219. doi: 10.1016/j.cell.2007.02.038.

Stoleru D, Peng Y, Agosto J and Rosbash M (2004) Coupled oscillators control morning and evening locomotor behaviour of Drosophila. Nature 2004 431:7010 431:862. doi: 10.1038/nature02926.

Vaze KM and Sharma VK (2013) On the Adaptive Significance of Circadian Clocks for Their Owners. Chronobiol Int 30:413–433. doi: 10.3109/07420528.2012.754457.

