## Supplemental Figure and Manual for "PHASE: A MATLAB Based Program for the Analysis of *Drosophila* Phase, Activity and Sleep under Entrainment"

*Running title: Software for Drosophila sleep and entrainment analysis*

[*]These authors contributed equally to this study.

**Supplementary Figure 1:** The effect of polynomial order (*p*) and window size (*n*) on the nature of fit between raw and smoothed data. Gray lines represent raw activity counts of flies over four consecutive cycles. Blue lines are smoothed data produced by the Savitzky-Golay filter.

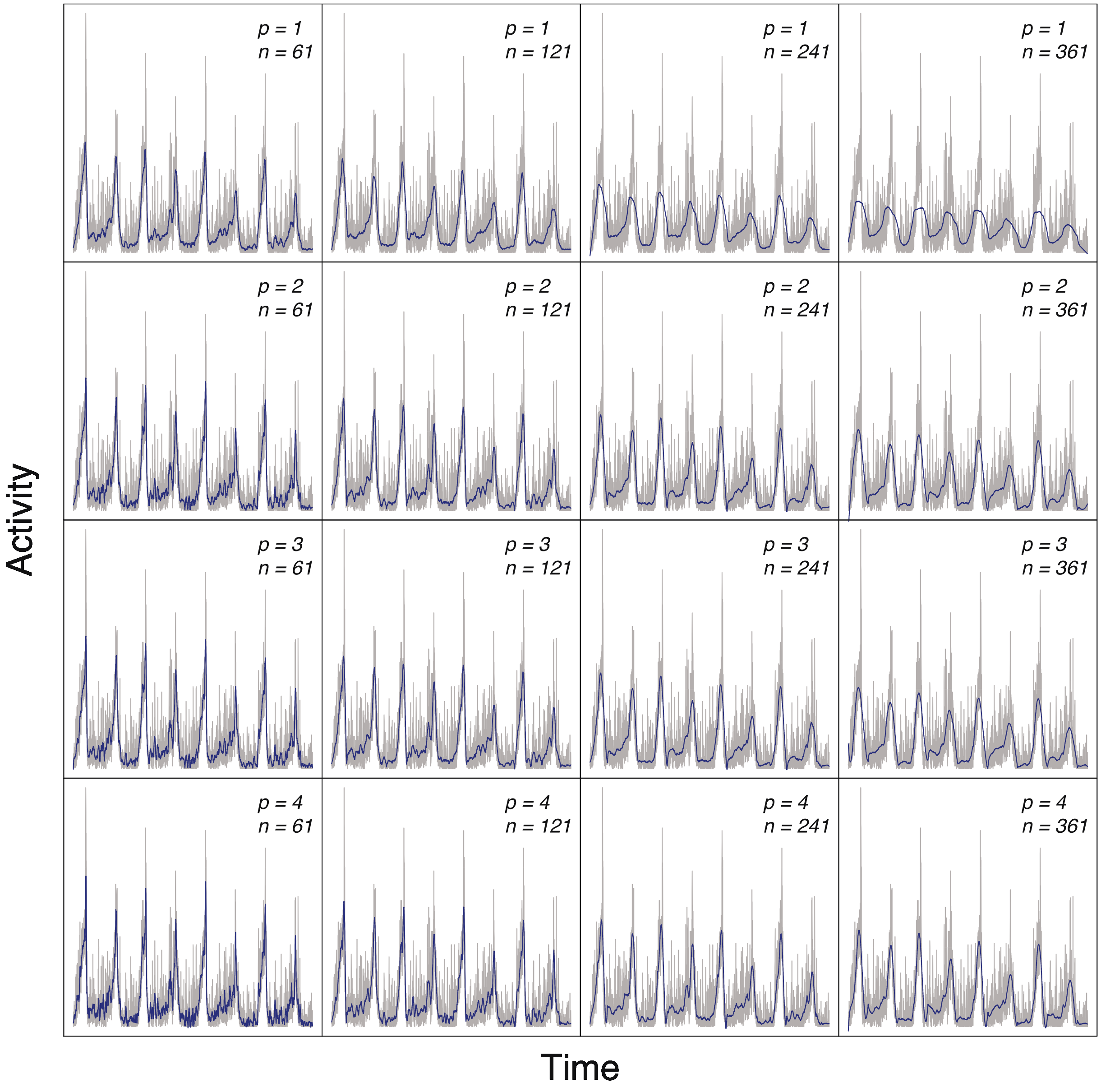

### PHASE (7.0) User Manual

### Table of Contents

***Section 1. Introduction ... 7***

***Section 2. PHASE Software ... 8***

**2.1. Software access ... 8**

**2.2. Getting started with PHASE ... 9**

**2.3. Example data sets ... 11**

**2.4. User-interface ... 11**

***Section 3. Input and raw data settings ... 12***

**3.1. Data processing protocol ... 12**

**3.2. Data settings ... 12**

***Section 4. Analysis types: data used, and input and output settings ... 15***

**4.1. Common export settings ... 15**

***Section 5. Profile analysis and visualization ... 16***

**5.1. Input ... 17**

**5.2. Output ... 19**

*5.2.1. Sleep* ... 19

*5.2.2. Normalized Activity* ... 21

*5.2.3. Averaged Activity* ... 23

**5.3. Graphs ... 25**

***Section 6. Phase analysis ... 26***

**6.1. Input ... 27**

**6.2. Output ... 28**

*6.2.1. Sleep* ... 28

*6.2.2. Normalized Activity* ... 29

*6.2.3. Averaged Activity* ... 30

### Section 1. Introduction

This manual is designed to guide users of PHASE in the analysis of *Drosophila* phase, activity, and sleep under entrainment, using data collected from the *Drosophila* Activity Monitor (DAM) system (TriKinetics Inc, Waltham, MA). Subsequent sections describe, (i) how to download, install and use the PHASE software, (ii) all PHASE analyses and the input settings required for each analysis, (iii) the Savitzky-Golay filtering used for several PHASE analyses, and (iv) all the output files generated and saved by PHASE.

PHASE may also prove useful for application outside of the *Drosophila* circadian field, provided that behavioral data is saved in a format that can be read and processed by TriKinetics *DAMFileScan* software.

***Note:*** *All figure citations in this manual are for figures described in the accompanying manuscript. Please refer to the manuscript for the figures.*

### Section 2. PHASE Software

#### 2.1. Software access

PHASE functions on both Mac (10.13 or later) and Windows (Windows 7 or later). There are two ways to run the PHASE software, (i) as a standalone program (a free and open-source version) or (ii) from within MATLAB (this requires a MATLAB license). To run PHASE as an App within MATLAB, users must acquire MATLAB Runtime R2020 (or later), the MATLAB Signal Processing Toolbox, which contains the Savitzky-Golay (SG) filtering function, and the MATLAB Financial Toolbox, which contains the user-interface for the calendar that is used in Data Settings of PHASE.

The standalone versions of PHASE for Mac OS and Windows may be downloaded from https://github.com/ajlopatkin/PHASE/tree/master/Installer%20Downloads (make sure to download operating system appropriate installer). See the example screenshot below to identify the button for downloading the installer(s). For further details please see the description in the README file or follow the instructions given below.

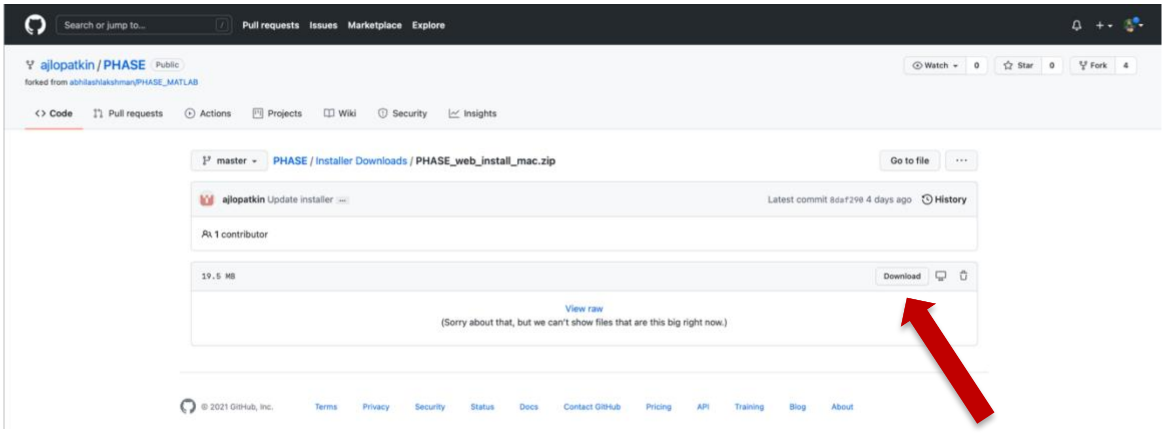

#### 2.2. Getting started with PHASE

1. Download the standalone bundle for your respective operating systems from here (see above image): https://github.com/ajlopatkin/PHASE/tree/master/Installer%20Downloads
2. Go to the folder containing the downloaded bundle and open the file.
3. Follow on-screen prompts. This is what your screen should look like when the download and installation have started:

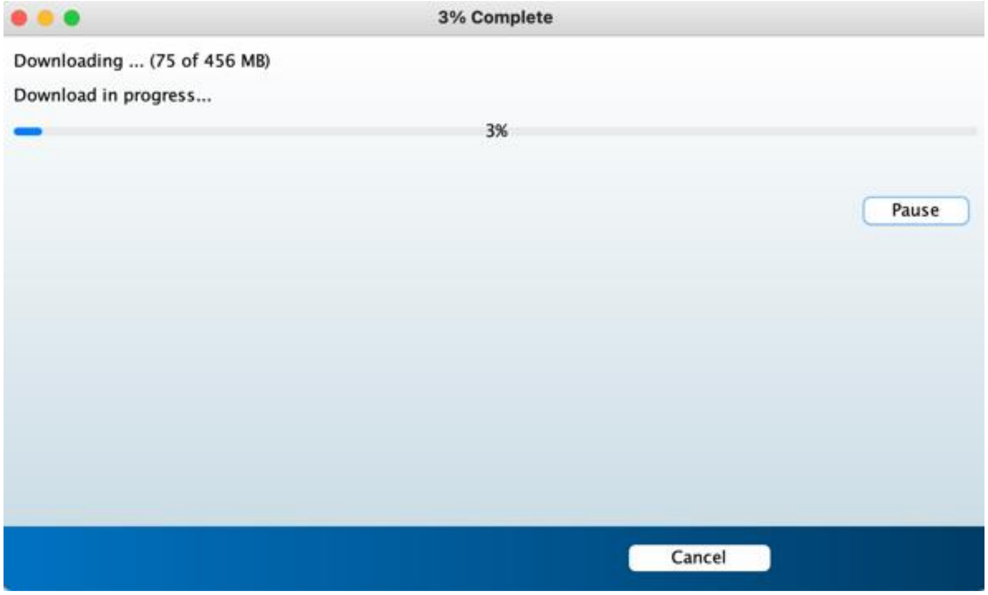

4. Go to your computer's applications folder (or the Desktop shortcut, in case of PC) and look for PHASEApp and launch it. The app should open and the following window should appear:

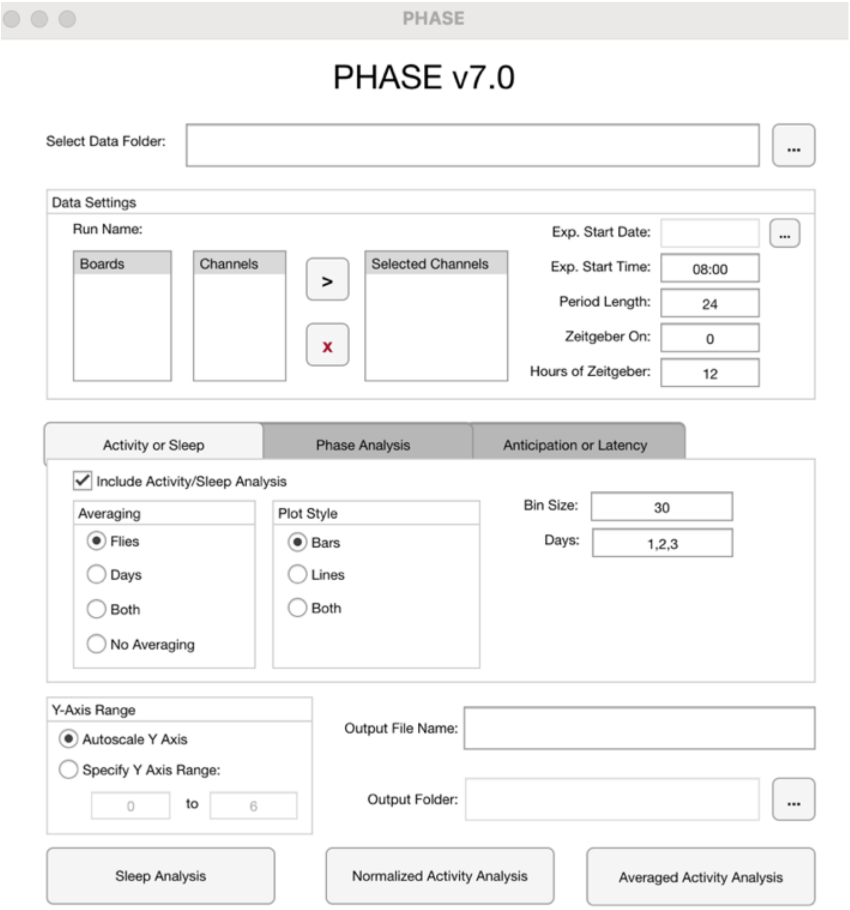

***Note:*** *Ensure that computer security settings allow downloads from the internet. MacOS Gatekeeper may disrupt PHASE file structure and successful installation.*

***Note:*** *To install and run PHASE via MATLAB, users must have MATAB Runtime R2020 or later. In addition, users will need to install the Signal Processing and Financial toolboxes. We recommend that users use the standalone versions because the app's behavior may be unexpected when run via MATLAB, depending on the MATLAB version and/or operating system.*

#### 2.3. Example data sets

The data sets used to demonstrate the utility of PHASE in the accompanying manuscript are available on GitHub (https://github.com/ajlopatkin/PHASE/tree/master/Example%20datasets). A guide to the data sets have been provided in Section 10.

#### 2.4. User-interface

The user-interface of PHASE can be segregated into five subsections (see Fig. 1 of accompanying manuscript). A detailed description of how to use the interface in subsequent sections and provided in the legend of the figure in the manuscript.

### Section 3. Input and raw data settings

#### 3.1. Data processing protocol

1. Use *DAMFileScan* (Trikinetics Inc, Waltham, MA) to process DAM monitor data into 1-minute bins and save them as channel files, for the entire duration of the experiment before analyzing it in PHASE.

   ***Note:*** *The 1-minute bin criterion is essential.*

#### 3.2. Data settings

1. Navigate to the folder containing DAM scanned channel files using the "…" button on the top right side (Fig. 1A-i).
2. Run Name, Exp. Start Date and Exp. Start Time should autofill. The Exp. Start Date and Exp. Start Time should be the exact same date and time of the first bin in the saved channel files.

   ***Note:*** *Data should be analyzed starting on the first bin, not on the start time. For example, for an experiment that started at 9PM (21:00) and was saved in 1-minute bins, the experimental start time should be set to 21:01.*

   ***Note:*** *Failure to autofill results may occur if there are channel files (from monitor files) with multiple start dates/times. In this case, users can simply use the calendar to choose relevant start dates/times.*

3. The left most box displays all the Monitors available in the chosen folder. The box on the right, next to the left most box displays all 32 channels of each monitor. PHASE can only analyze channel files from one monitor at a time.
4. Highlight the channels from the second box to chose the individual flies that must be included in the analysis. To select multiple channels, hold control or command and click the desired channels.
5. Use the black ">" arrow to push the selected channel files to the analysis box (third box). Any channel listed on the third box will be included in the analysis.
6. Use the red "X" to clear all channel files before loading new channel files for analysis.

***Note:*** *If incorrect channel files were selected by accident, use the red "X" button to clear all the channel files and select relevant channel files once again.*

7. Change auto-filled "Exp. Start Date" using the "…" button from the right side of the interface, to select the date on which analysis must begin.
8. Change the auto-filled "Exp. Start Time" to the local time corresponding to the time at which the analysis will start in HH:MM format. For instance, if lights turned on in an LD cycle at 10AM, and users would like to start the analysis at ZT00, then the corresponding Exp. Start Time would be 10:01.

   ***TIP:*** *For appropriate analysis of peaks and certain aspects of anticipation, it is critical that the two peaks of activity are centered with respect to the duration of the day such that none of the expected peaks are at the start or end of the cycle being analyzed. If the peaks occur at the edges, estimates for those peaks will be inaccurate. For instance, if users expect peaks around ZT00 and ZT12 and the local time of lights-ON (for an LD 12:12 cycle) is at 10AM, an appropriate Exp. Start Time would be 4:01AM and the corresponding Zeitgeber ON would be 6 (i.e., six hours after the experimental start time). Also see Section 11 for reproducible examples.*

9. Change "Period Length", "Zeitgeber ON", and "Hours of Zeitgeber" as needed.
10. "Period Length" is the duration of one environmental cycle in hours.
11. "Zeitgeber ON" refers to the number of hours after the Exp. Start Time that the zeitgeber turns ON. For instance, if lights turn on at 10AM, and the Exp. Start Time was set as 10:01, then "Zeitgeber ON" must be 0. If the Exp. Start Time was set as 4:01, then "Zeitgeber ON" must be 6, because zeitgeber turned ON 6-h after the Exp. Start Time, and so on.
12. "Hours of Zeitgeber" refers to the duration of the entire cycle that the zeitgeber was in the ON state, in hours. For example, if flies were entrained to a 12:12 LD cycle, the appropriate "Hours of Zeitgeber" would be 12.

    ***Note:*** *This value can be changed in order to perform window-specific analyses, as will be demonstrated using reproducible examples in Section 11 (Also see Fig. 3 in the accompanying manuscript).*

13. Auto-fills for “Period Length”, “Zeitgeber ON” and “Hours of Zeitgeber” are “24”, “0” and “12”, respectively.

    ***Note:*** *“Day” statistics are derived from the hours between “Zeitgeber ON” and OFF (OFF times are estimated using the “Hours of Zeitgeber”). “Night” statistics are derived from the remaining hours of the cycle being analyzed.*

### Section 4. Analysis types: data used, and input and output settings

PHASE can visualize and both activity and sleep profiles, objectively quantify the phases of behavioral peaks, and estimate anticipation or latency of activity or sleep behavior. To analyze sleep data users must click the "Sleep Analysis" on the bottom left side (Fig. 1A-v). To analyze normalized or averaged activity data, users must click the "Normalized Activity Analysis" or "Averaged Activity Analysis" buttons in the middle and right side, respectively (Fig. 1A-v).

PHASE bins the raw data into user defined intervals, can average activity/sleep across cycles, across individuals, or across both cycles and individuals, exports all this data as a comma separated file which can be opened using MS-Excel, and plots these data and exports these figures as MATLAB compatible figures and PDF files. Furthermore, PHASE uses a Savitzky-Golay (SG) filter to smooth raw data, objectively define phases and computes measures of sleep latency, and activity anticipation. Each of these analyses requires specific inputs, which are described in the following sub-sections.

#### 4.1. Common export settings

Before providing detailed guidance for each of the analysis methods mentioned above, we will introduce the users to section iv of Fig. 1, which determines the *y*-axis scale of output figures and the name and location of the files produced by PHASE analysis.

1. Users must choose *y*-axis range to appear in the output plots of PHASE. Users can either allow PHASE to automatically adjust *y*-axis or can specify the *y*-axis range (Fig. 1A-iv, left).
2. Users must also select an "Output Folder" where all files and plots will be saved.
3. Users must also provide an "Output File Name" that uniquely identifies the set of analyses run by PHASE.

***TIP:*** *Create a new folder for PHASE's output files.*

### Section 5. Profile analysis and visualization

The sleep/activity analyses on PHASE requires the following inputs. To perform these analyses, check the checkbox “Include Activity/Sleep Analysis” under the “Activity or Sleep” tab (Fig. 1A-iii).

***Note:*** *A single bout of “sleep” is defined as five consecutive 1-minute bins (or longer) with zero infrared beam crossings (Hendricks et al., 2000; Huber et al., 2004). PHASE quantifies sleep this way.*

#### 5.1. Input

1. Indicate if PHASE will analyze activity/sleep and visualize profiles by averaging across “Flies,” “Days,” or “Both” or if “No Averaging” must be done. Outputs produced by each of these options is described below:
    a. Averaging by “Flies” averages behavior of all selected flies across all the chosen “Days” and provides a single graph with each day of data displayed consecutively. In other words, the averaged timeseries of activity/sleep is plotted.
    b. Averaging by “Days” averages across all the chosen “Days” for each selected fly. This will produce a single full-cycle plot of activity/sleep for each chosen fly.
    c. “Both” results in a single plot that averages activity/sleep, first across all chosen “Days” and then across all chosen “Flies.”
    d. “No Averaging” creates one timeseries for every fly displaying the raw data across all the chosen “Days”.
2. Indicate the desired nature of profile plots. The user can can choose to visualize the data as bar graphs, line plots, or both.
    a. “Bars” creates a traditional vertical bar graph displaying binned activity or sleep data with standard error of mean (SEM) error bars.
    b. “Lines” creates a line graph with SEM shaded above and below the mean line.
    c. “Both” creates both types of graphs.

***Note:*** *The standard errors displayed by PHASE are the standard errors across flies, not across days.*

3. Indicate the desired binning interval in the output by entering value in the “Bin Size” box.

***Note:*** *The bin size must be in minutes. For instance, bin size of 1-h must be input as 60.*

4. Indicate the days of analysis that PHASE will include for sleep analysis. Multiple days can be input here, with each day separated by a comma (for instance, 1,2,3). The days don’t necessarily have to be consecutive. For example, 1,2,3,5 is a valid input.

***Note:*** *Day "1" begins at the "Exp. Start Time" for the indicated "Day Length" with "Day" beginning at "Zeitgeber ON" and "Night" beginning after the given "Hours of Zeitgeber" in "Data Settings". Day "2" begins after the indicated "Day Length" following the start of day "1".*

#### 5.2. Output

##### *5.2.1. Sleep*

When Sleep Analysis is executed (the button shown in Fig. 1A-v), MS-Excel compatible file(s) is/are generated with the name: "SleepAnalysis_OutputFileName_YYYMMDD_HHMM". The following is a description of the sheets that are created by PHASE in the MS-Excel file(s).

1. SummedStatistics (Raw)

   This sheet contains the following columns of data, values of which are summed over all indicated days in the "Days" input.

   A. *Board:* Monitor number (1-120)

   B. *Fly:* Channel number (1-32)

   C. *TotalMinutes:* Sum of all data intervals with non-zero sleep values multiplied by the length of that interval (in minutes).

   D. *AverageMinutes:* The "TotalMinutes" divided by the total number of data collection bins.

   E. *TotalBouts:* The total number of bouts of sleep. A bout is defined as a series of non-zero values of any length that qualifies as sleep.

   F. *AverageBoutDuration:* The sum of the bout durations in minutes divided by the total number of bouts.

   G. *DayTotal:* Total minutes of sleep during "Zeitgeber ON" hours.

   H. *AverageDayMinutes:* The "DayTotal" divided by the total number of data collection bins in duration between "Zeitgeber ON" and Zeitgeber OFF.

   I. *DayBouts:* The total number of bouts beginning during "Zeitgeber ON" hours.

   J. *AverageDayBoutDuration:* The total minutes of sleep bouts that began during "Zeitgeber ON" hours divided by the total number of bouts that during the same time.

   K. *NightTotal:* Total minutes of sleep outside the "Zeitgeber ON" hours.

   L. *AverageNightMinutes:* The "NightTotal" divided by the total number of data collection bins outside of the "Zeitgeber ON" hours.

   M. *NightBouts:* The total number of bouts beginning outside the "Zeitgeber ON" hours.

N. *AverageNightBoutDuration:* The total minutes of sleep bouts that began after "Zeitgeber ON" hours divided by the total number of bouts that during the same time.

2. DayAvgStatistics (Raw)

   This sheet contains all the column-wise information as described for the SummedStatistics sheet above, but averaged across cycles, instead of sum.

3. BinnedData (Raw)

   Contents of this sheet will depend on what averaging option was chosen, i.e., "Flies", "Days", "Both" or "No Averaging."

   a. When averaged across "Flies", data are presented as time-series averaged over all individuals.
   b. When averaging across "Days", each subsequent column will have individual fly sleep data averaged across all the days. When PHASE reports individual fly data, it stores the identity by using the monitor number and channel number. For instance, channel 2 of monitor 13, will be represented as M013C02.
   c. When averaging across "Both", flies and days, a single column of data is presented.
   d. When "No Averaging" is selected, raw time series through the selected days for every fly is reported.

***Note:*** *Time is referred to as t_Xmin. The first time-point (based on the Exp. Start Time) is referred to as t_0min, with every subsequent bin being labeled so forth. For instance, the second time-point in case the user chooses 30-min bins would be t_30min.*

4. Settings

   This sheet records all the settings chosen by the using for the analysis session.

5. Daily data sheets

   All other sheets record sleep statistics and sleep profiles for each day subjected to analysis, as described above, and results for each day are save ine separate files. For instance, if users chose Days 1,3,4 for analysis these sheets will be called, "Day 1 Statistics (Raw)", "Day 1 BinnedData (Raw)", "Day 3 Statistics (Raw)", "Day 3 BinnedData (Raw)," and so on.

##### 5.2.2. *Normalized Activity*

Exports from "Normalized Activity Analysis" are summarized in the multi-sheet MS-Excel file: "NormalizedActivityAnalysis_OutputFileName_YYYMMDD_HHMM". The following is a description of the sheets that are created by PHASE.

***Note:*** *Normalization is done relative to total activity counts in one user-defined cycle.*

1. SummedStatistics (Raw)

   This sheet contains the following columns of data, values of which are summed over all indicated days in the "Days" input.

   A. *Board:* Monitor number (1-120)
   B. *Fly:* Channel number (1-32)
   C. *TotalCounts:* The sum of all counts of activity in the data series for the selected days.
   D. *AverageCounts:* The "TotalCounts" divided by the total number of bins. E.g., for 1-min scanned files, analyzed over three 24-hour days would divide all "TotalCounts" values by (1440×3).
   E. *DayTotalCounts:* The total IR beam crosses that took place during "Zeitgeber ON" hours.
   F. *AverageDayCounts:* The "DayTotalCounts" divided by the total number of data collection bins in "Zeitgeber ON" hours.
   G. *NightTotalCounts:* The total IR beam crosses that took place outside of the "Zeitgeber ON" hours.
   H. *AverageNightCounts:* The "NightTotalCounts" divided by the total number of data collection bins outside of "Zeitgeber ON" hours.

2. DayAvgStatistics (Raw)

   This sheet contains all the column-wise information as described above, but averaged over cycles, instead of summed.

3. BinnedData (Raw)

   Contents of this sheet will depend on what averaging option was chosen.

a. When averaged over "Flies," data are presented as a timeseries averaged over all individuals.
b. When averaging over "Days," each subsequent column will display individual fly activity data averaged across all days. When PHASE reports individual fly data, it indicates the identity of each fly using the monitor number and channel number. For instance, channel 2 of monitor 13, will be represented as M013C02.
c. When averaging across "Both", flies and days, a single column of data is presented.
d. When "No Averaging" is selected, raw timeseries across the selected days for every fly is reported.

4. SummedStatistics

   This sheet contains the same contents as that in SummedStatistics (Raw) but using normalized data. For instance, TotalCounts normalized for 3 days of analysis for every fly will be equal to 3. This is because activity is normalized to total activity over a cycle, which must therefore add up to one for each day.

5. DayAvgStatistics

   This sheet contains the same contents as that in DayAvgStatistics (Raw) but using normalized data as described above.

6. BinnedData

   Contents of this sheet are in the same format as that in the sheet BinnedData (Raw). This contains activity counts normalized to total counts of activity within that day.

7. Settings

   This sheet records all the settings utilized for the specific analysis session.

8. Daily data sheets

   All other sheets record activity statistics for each analyzed day and activity profiles in the same way described above, but for each cycle separately. For instance, if users chose Days 1,3,4 for analysis these sheets will be called, "Day 1 Statistics (Raw)", "Day 1 BinnedData (Raw)", "Day 1 Statistics", "Day 1 BinnedData", "Day 3 Statistics (Raw)", "Day 3 BinnedData (Raw)", "Day 3 Statistics", "Day 3 BinnedData" and so on.

*5.2.3. Averaged Activity*

Exports from "Averaged Activity Analysis" are summarized in the multi-sheet MS-Excel file: "AveragedActivityAnalysis_OutputFileName_YYYMMDD_HHMM". The following is a description of the sheets that are created by PHASE.

***Note:*** *Averaged activity refers to mean number of beam crossings within each user-defined Bin.*

1. SummedStatistics (Raw)
   Contents of this sheet are the same as described under the Normalized Activity section.
2. DayAvgStatistics (Raw)
   Contents of this sheet are the same as described under the Normalized Activity section.
3. BinnedData (Raw)
   Contents of this sheet are the same as described under the Normalized Activity section.
4. SummedStatistics
   This sheet contains the same contents as that in SummedStatistics (Raw) but using averaged data.
5. DayAvgStatistics
   This sheet contains the same contents as that in DayAvgStatistics (Raw) but using averaged data.
6. BinnedData
   Contents of this sheet are in the same format as that in the sheet BinnedData (Raw). This contains total activity counts in the user-defined bin divided by the user-defined bin size. In other words, values represent the average activity in a given bin.
7. Settings
   This sheet records all the settings utilized for the specific analysis session.
8. Daily data sheets
   All other sheets record day-wise activity statistics and activity profiles in the same way described above, but for each cycle separately. For instance, if users chose Days 1,3,4 for analysis these sheets will be called, "Day 1 Statistics (Raw)", "Day 1 BinnedData (Raw)",

“Day 1 Statistics”, “Day 1 BinnedData”, “Day 3 Statistics (Raw)”, “Day 3 BinnedData (Raw)”, “Day 3 Statistics”, “Day 3 BinnedData” and so on.

#### 5.3. Graphs

Graphs generated by PHASE represent activity or sleep data collected in the "Bin Size" specified for each individual fly in "Data Settings". When sleep is analyzed, the *y*-values on the graph represent total minutes of sleep in the user-defined bin. When normalized activity is analyzed, the *y*-values represent activity normalized to total activity in the entire cycle. When averaged activity is analyzed, the *y*-values represent activity in each user-defined bin averaged for the user-defined bin size. Bar and line graphs represent Zeitgeber OFF hours with darker bars or shaded background regions, respectively. Bar graphs display SEM as error bars above the mean activity/sleep bar. Line graphs shade the SEM area above and below a darker, mean activity/sleep line.

#### Section 6. Phase analysis

“Phase Analysis” applies a Savitzky-Golay filter (see Section 9 and the accompanying manuscript for details on this filter and how it is used) to sleep, normalized or averaged activity depending on the analysis chosen by the user.

***Note:*** *PHASE only detects phases of peaks, and not other circadian phase markers such as onset, offset and trough.*

#### 6.1. Input

1. Indicate that PHASE must analyze phases by checking the box labelled "Include Phase Analysis" (see Fig. 1B).
2. Indicate the "Filter Order" to be used (see main text and Sections 9 and 11 for clarity on "Filter Order").
3. Indicate the "Filter Frame Length (minutes)" to be used (see Sections 9 and 11 for clarity on "Filter Frame Length").

***Note:*** *Filter frame length must be an odd number.*

4. Indicate the ZT around which phases must be called. Multiple ZT values can be used separated by commas.
5. Indicate what the "Minimum Distance Between Peaks" must be. This value must be provided in minutes. PHASE uses the MATLAB function *findpeaks* to find local maxima outside the "Minimum Distance Between Peaks" of filtered data. The "Minimum Distance" restricts the *findpeaks* function to an acceptable peak-to-peak distance (in minutes), effectively ignoring any peaks which are close together. Therefore, smaller minimum distances will allow PHASE to find many, small peaks. Larger minimum distances will favor larger, dominant peaks.
6. Indicate which days will be used to identify phases. Multiple days can be used as input, separated by commas. Where multiple days are specified in "Days" averaging across days takes place prior to filter application.

***Note:*** *If phase calls for single days are required, users must analyze one day at a time.*

#### 6.2. Output

##### *6.2.1. Sleep*

Sleep peak phases are exported as: "SleepPhase_OutputFileName_YYYMMDD_HHMM". This is a two-sheet file which have the following contents:

1. Phase

   This sheet contains three columns for every ZT chosen in the phase analysis input section. The values for each ZT are in descending order of the ZT value. For instance, if two ZT values are used in the input, i.e., 0,12, columns B, C and D will report Peak Time (ZT), Peak Height and Peak Area, respectively around ZT12. Columns E, F and G will report the same for ZT0, and so on. Column A will always be the monitor and channel identity of every fly in the format M*XXX*C*YY*, such that a fly from monitor 1, channel 3 will be labeled as M001C03.

   *Peak Time (ZT):* PHASE picks the time (ZT) at which the maximum sleep occurs. ZT is computed relative to the provided input in the "Data Settings".

   *Peak Height:* PHASE uses the function *findpeaks* to also determine the peak height/prominence (in minutes of sleep) and the peak width/half-prominence (in minutes). Prominence of a peak is a measure of how the peak stands out in relation to its height and location compared to other neighboring peaks (see https://www.mathworks.com/help/signal/ug/prominence.html for more details).

   *Peak Area:* This is calculated as the product of Peak Height and Peak Width.

***Note:*** *PHASE can find, quantify and visualize all peaks within smoothed/filtered behavior data. However, it is advisable that users search for peaks closest to a given ZT, or set of ZT(s).*

2. Settings

   This sheet records all the settings utilized for the specific analysis session.

##### *6.2.2. Normalized Activity*

Phase calls on 1-minute binned normalized (by total daily activity) activity data are summarized in the two-sheet Excel file "NormalizedActivityPhase_OutputFileName_YYYMMDD_HHMM".

1. Phase
   Contents of this sheet are in the same format as described for the output of sleep data as described above.
2. Settings
   This sheet records all the settings utilized for the specific analysis session.

##### *6.2.3. Averaged Activity*

Phase calls on averaged activity data are summarized in the two-sheet Excel file "ActivityPhase_OutputFileName_YYYMMDD_HHMM".

1. Phase

   Contents of this sheet are in the exact same format as described for the output of sleep data as described above.
2. Settings

   This sheet records all the settings utilized for the specific analysis session.

#### 6.3. Graphs

Graphs generated from this analysis represent sleep, normalized or averaged activity for each individual fly included in “Data Settings” for the day or average of days indicated in “Days”. Blue lines represent the filtered data curve of the 1-minute data. Graphs indicate peak areas in blue shading and label the peak maximum with a blue arrowheads and ZT values (Figures 4A and 5A).

### Section 7. Anticipation or Latency analysis

"Anticipation or Latency Analysis" applies an SG filter (see Section 9 and accompanying manuscript for details on this filter and how it is used) to sleep/activity data.

***Note:*** *The slope method of activity anticipation is performed using raw binned data only when Averaged Activity Analysis is chosen by the user.*

#### 7.1. Input

1. Perform anticipation or latency analysis by checking the box labelled “Include Anticipation or Latency Analysis”.
2. Indicate the “Filter Order” to be used (see Section 9, accompanying manuscript and Suppl. Fig. 1 for clarity on “Filter Order”).
3. Indicate the “Filter Frame Length (minutes)” to be used (see Section 9, accompanying manuscript and Suppl. Fig. 1 for clarity on “Filter Frame Length”).

***Note:*** *Filter frame length must be an odd number.*

4. Indicate “Window Length (minutes)” to be used. All anticipation and/or latency analyses are performed within this user-defined window length.
5. Indicate the ZT around which phases will be identified. Multiple ZT values can be used separated by commas.
6. Indicate which days will be analyzed for anticipation and/or latency. Multiple days can be used as input, separated by commas. When multiple days are specified in “Days” averaging across days takes place prior to filter application.

***Note:*** *If single day data for anticipation and/or latency are required, users must analyze time-series data one day at a time.*

#### 7.2. Output

##### *7.2.1. Sleep*

Data are summarized in the file: "SleepLatency_OutputFileName_YYYMMDD_HHMM". Whenever multiple days are chosen for analysis, averaging is performed before filtering or calculating slopes. Analysis of latency around each user-defined ZT is provided in a separate Excel sheet. The columns in each of these sheets contain the following data in the following order.

A. The left-most column has monitor and channel identity of every fly in the format M*XXX*C*YY*, such that a fly from monitor 1, channel 3 will be labeled as M001C03.

B. *Min Sleep Time (ZT):* PHASE moves positively along the *x*-axis (time) from a user-defined ZT to derive the first minima/least amount of sleep from filtered data. The time (ZT) at which this occurs is reported in this column.

C. *Max Sleep Time (ZT):* PHASE moves positively along the *x*-axis from a user-defined ZT to derive the maxima/greatest amount of sleep from filtered data. This time value (ZT) is reported in this column.

D. *Latency (minutes):* PHASE determines the duration in minutes between defined maxima and minima and reports this latency value for each fly in this column.

E. *Index (AUC):* This column documents area under the curve (AUC) of the filtered data between the minima and maxima defined above, for every fly.

F. *Slope:* PHASE also finds the slope of a linear regression fitted to unsmoothed, single fly sleep data for each user-defined ZT within the indicated "Window Length (minutes)".

##### *7.2.2. Normalized Activity*

Anticipation analysis performed on SG filtered 1-minute binned normalized activity data are summarized in: “NormalizedActivityAnticipation_OutputFileName_YYYMMDD_HHMM”. Similar to the case with sleep, whenever multiple days are chosen for analysis, averaging is performed before filtering or calculating slopes. Analysis of anticipation around each user-defined ZT is provided in a separate Excel sheet. The columns in each of these sheets contain the following data in the following order.

A. The left-most column has monitor and channel identity of every fly in the format M*XXX*C*YY*, such that a fly from monitor 1, channel 3 will be labeled as M001C03.
B. *Min Activity Time (ZT):* PHASE moves negatively along the *x*-axis (time) from a user-defined ZT to derive the first minima/least amount of activity from filtered data. The time (ZT) at which this occurs is reported in this column.
C. *Max Activity Time (ZT):* PHASE moves negatively along the *x*-axis from a user-defined ZT to derive the maxima/greatest amount of activity from filtered data. This time value (ZT) is reported in this column.
D. *Anticipation (minutes):* PHASE determines the duration in minutes between defined maxima and minima and reports this as the duration of anticipation for each fly in this column.
E. *Index (AUC):* This column documents area under the curve (AUC) of the filtered data between the minima and maxima defined above, for every fly.
F. *Slope:* PHASE also finds the slope of a linear regression fitted to unsmoothed normalized activity data for individual flies for each user-defined ZT within the indicated “Window Length (minutes)”.

##### *7.2.3. Averaged Activity*

Anticipation analysis performed on SG filtered 1-minute binned averaged activity data are summarized in: “ActivityAnticipation_OutputFileName_YYYMMDD_HHMM”. Whenever multiple days are chosen for analysis, averaging is performed before filtering or calculating slopes. Analysis of anticipation around each user-defined ZT is provided in a separate Excel sheet. The columns in each of these sheets contain the same contents as those described above for the Normalized Activity section, except *Slope*.

For *Slope* analyses, PHASE uses raw binned data to fit linear regressions within the user-defined window and extracts estimates of slope.

#### 7.3. Graphs

##### *7.3.1. Index and Duration*

Graphs represent 1-minute binned raw activity data (when “Averaged Activity Analysis” is selected), normalized activity data (“Normalized Activity Analysis”), or sleep data for each individual fly included in “Data Settings” for the day or average of days indicated in “Days”. Each region included in the analysis is shaded in light grey. The blue line represents the filtered data curve. Blue triangles indicate the filtered activity or sleep maximum and minimum bins determined by PHASE within the “Window Length” from each ZT. The time (in minutes) between each maximum and minimum is annotated for each region above the blue arrowhead (Fig. 6D).

##### *7.3.2. Slope*

Graphs represent slopes calculated from unsmoothed binned activity data (“Averaged Activity Analysis”), unsmoothed normalized activity data (“Normalized Activity Analysis”) or sleep data within the analysis window of a given ZT for all flies included in “Data Settings” (Figs. 6A and B).

As sleep values range from 1 or 0 for a given bin, these values are far more discreet than raw activity values (presented as unsmoothed IR beam crosses). For example, given the 0-or-1 sleep values, for three days of activity there are only four possible average values: 0, 1/3, 2/3, or 1 depending how many of those days the fly was asleep during each interval.

In either case, sleep/normalized activity/averaged activity, regression plots for each ZT value included in the “Anticipation or Latency Analysis” is plotted separately. The linear regression/slope for individual flies is in light grey. The average linear regression/slope for all flies included in analysis is in dark grey (Figs. 6A and B).

#### Section 8. Graph tools

Once a graph has been generated by PHASE, a window containing it will automatically open. Clicking on the graph will display a number of graph tools on the top of the graph. A description of these tools follows.

1. House: Restore original graph, undo any changes.
2. Magnifying glasses: Zoom in and out.
3. Hand: Move the graph.
4. Data tip: Gives exact ($x$,$y$) coordinates of selected data point.
   To remove data tips, right click the data tip box and select 'remove data tip'.
5. Paintbrush: Highlight specific data, e.g., a bar.
6. Arrow with box: Save or copy graph.
   ***Note:*** *To further edit using MATLAB copy as vector graphic. For a fixed image of the graph copy as image.*

7. Legend: In the toolbar above the graph, select the white box with red and blue squares. Customize by right clicking.

### Section 9. Savitzky-Golay Filter

PHASE uses the MATLAB function *sgolay* from the Signal Processing Toolbox to filter 1-minute activity or sleep data with a Savitzky-Golay finite impulse response (FIR) smoothing filter of a given "Filter Order" and "Filter Frame Length". Savitzky-Golay filtering is optimal for smoothing data with a large frequency span (such as 1-minute activity data); reducing signal-to-noise while preserving the original shape of signals that are often distorted using standard averaging filters (Savitzky and Golay, 1964; Schafer, 2011). The function *sgolay* replaces successive central points in frames (minutes of behavior within the "Filter Frame Length") with the weighted average of all measures within the frame. The weighted coefficients are obtained by fitting all frame values to a polynomial of a given degree ("Filter Order") using the linear least squares method.

PHASE applies a low-order polynomial function specified by the "Filter Order" across an odd-number of minutes of activity or sleep behavior within the "Filter Frame Length". The "Filter Order" and "Filter Frame Length" is user-specified in "Phase Analysis", and "Anticipation or Latency Analysis". Therefore, it is imperative that users find the best representation of activity and sleep data that minimizes noise and signal distortion and maximizes signal. This is an iterative process until the final polynomial order and frame length are defined. To best utilize PHASE's smoothing functions, users may begin by generating activity or sleep graphs using the autofilled parameters for "Filter Order" and "Filter Frame Length". After visual inspection, users should apply the following principles of Savitzky-Golay filtering in order to more accurately represent their data and avoid generating erroneous estimates of phase and anticipation and/or latency. Please refer to the accompanying manuscript, Suppl. Fig. 1 and the reproducible examples in the next section, to get a better idea of how to use the SG filter.

***Note:*** *The polynomial degree in "Filter Order" must not exceed the minutes defined by "Filter Frame Length".*

***Note:*** *The "Filter Frame Length" must be an odd number. SG methodology assumes that the data within the specified frame are equidistant. Symmetry requires the window to contain an odd number of points in order to replace a central point by the best representation of all values within the frame.*

***Note:*** *Signal-to-noise generally increases as the polynomial degree decreases. A higher "Filter Order" will best preserve narrow feature height and width but perform less well on broad peaks. A lower "Filter Order" conversely, will smooth the data reducing time-series to their broad features.*

***Note:*** *Signal-to-noise increases as "Filter Frame Length" increases as there are more data points over which to derive the weighted average of the center point. Increased frame length also increases original signal distortion, however.*

***Note:*** *The ratio between the "Filter Order" and "Filter Frame Length" controls original data distortion. Smoothing generally increases as the ratio between the two values increases.*

#### Section 10. Guide to sample data sets

Two sets of scanned channel files are provided as example data sets (https://github.com/ajlopatkin/PHASE/tree/master/Example%20datasets). These are the same data sets used to generate all the analyses reported in the accompanying manuscript.

***Set-1:*** These channel files are named "ScannedCtMxxxCyy.txt." The "xxx" refers to the monitor number and the "yy" refers to the channel number. All the data in this set are used in Figs. 2-6, except Figs. 3G and H.

| **Monitor Number** | **Channels** | **Genotype and sex** | **Experimental regime** | **Exp. Start Date** | **Lights-ON** |
|---|---|---|---|---|---|
| 23 | 1-32 | *Canton-S*, males | LD12:12 | 20 Dec 2020 | 21:00 |
| 59 | 1-32 | *Canton-S*, males | LD08:16 | 20 Dec 2020 | 21:00 |
| 67 | 1-32 | *Canton-S*, males | LD16:08 | 20 Dec 2020 | 21:00 |
| 56 | 1-32 | $per^{01}$, males | LD12:12 | 20 Dec 2020 | 21:00 |
| 64 | 1-32 | $per^{01}$, males | LD08:16 | 20 Dec 2020 | 21:00 |
| 72 | 1-32 | $per^{01}$, males | LD16:08 | 20 Dec 2020 | 21:00 |
| 47 | 1-32 | $w^{1118}$, males | LD12:12 | 25 Jun 2021 | 11:00 |
| 48 | 1-32 | $pdf^{01}$, males | LD12:12 | 25 Jun 2021 | 11:00 |

***Set-2:*** These channel files are named "ChannelsCtMxxCyy.txt." The "xxx" refers to the monitor number and the "yy" refers to the channel number. All the data in this set are used only in Figs. 3G and H.

| **Monitor Number** | **Channels** | **Genotype and sex** | **Experimental regime** | **Exp. Start Date** | **Lights-ON** |
|---|---|---|---|---|---|
| 22 | 1-32 | *Canton-S*, males | LD12:12 | 10 Apr 2021 | 09:00 |
| 23 | 1-32 | *Canton-S*, females | LD12:12 | 10 Apr 2021 | 09:00 |

#### Section 11.  Reproducing results in the accompanying manuscript

All the data used in the accompanying manuscript to illustrate the use of PHASE are freely available on GitHub (link for download in the previous section).  All the data are segregated into two sets, i.e., *Set-1* and *Set-2*.  *Set-1* has the raw data files for all figures except for Figs. 3G and 3H.  Raw data for the latter two figures are in *Set-2*.

Before attempting to reproduce the following analyses, we recommend the user to download and extract all the raw data to a local folder.

#### 11.1. Replicating Fig. 2

1. Click the "…" button as shown in Fig. 1A-i and navigate to the folder containing the data *Set-1*.
2. Select all 32 channels of Monitor 23.
3. Set Exp. Start Date to 20-Dec-2020, Exp. Start Time to 21:01, Period Length to 24, Zeitgeber On to 0, and Hours of Zeitgeber to 12.
4. Check the "Include Activity/Sleep Analysis" button (Fig. 1A-iii).
5. Choose Averaging as "Days" and Plot Style as "Lines". Set Bin Size to 30 and Days to 1,2,3,4,5,6,7.
6. Type an output file name and navigate to any desired output folder location.
7. Click the "Averaged Activity Analysis" button (Fig. 1-A-v).
8. All relevant figures and data for Fig. 2A should have been generated.
9. Repeat the same steps as above but choose averaging options of "Flies" and "Both" to generate relevant plots and data for Figs. 2B and C, respectively.

##### 11.2. Replicating Fig. 3

1. Click the "…" button as shown in Fig. 1A-i and navigate to the folder containing the data *Set-1*.
2. Select all 32 channels of Monitor 23.
3. Set Exp. Start Date to 20-Dec-2020, Exp. Start Time to 21:01, Period Length to 24, Zeitgeber On to 0, and Hours of Zeitgeber to 12.
4. Check the "Include Activity/Sleep Analysis" button (Fig. 1A-iii).
5. Choose Averaging as "Days" and Plot Style as "Lines". Set Bin Size to 30 and Days to 1,2,3,4,5,6,7.
6. Type an output file name and navigate to any desired output folder location.
7. Click the "Sleep Analysis" button (Fig. 1-A-v).
8. All relevant figures and data for Fig. 3A should have been generated.
9. Repeat the same steps as above but choose averaging options of "Flies" and "Both" to generate relevant plots and data for Figs. 3B and C, respectively.

10. Click the "…" button as shown in Fig. 1A-i and navigate to the folder containing the data *Set-2*.
11. Select all 32 channels of Monitor 22.
12. Set Exp. Start Date to 10-Apr-2021, Exp. Start Time to 14:01, Period Length to 24, Zeitgeber On to 0, and Hours of Zeitgeber to 3.

    ***Note:*** *In this experiment, lights ON was at 9:00AM which means that 14:01 is the start of ZT05. By setting the Hours of Zeitgeber to 3, we instruct PHASE to consider the three-hour window starting at ZT5 to be "daytime". Therefore, all activity/sleep statistics recorded as happening during the daytime is happening between ZT05-08. In this way, users can get window specific activity/sleep statistics of interest.*

13. Check the "Include Activity/Sleep Analysis" button (Fig. 1A-iii).
14. Choose Averaging as "Flies" and Plot Style as "Lines". Set Bin Size to 30 and Days to 1,2,3.
15. Type an output file name and navigate to any desired output folder location.

16. Click the “Sleep Analysis” button (Fig. 1-A-v).
17. All relevant figures and data for Fig. 3G-top should have been generated. This is the sleep timeseries for male flies.
18. Repeat steps 10-16 but select all 32 channels of Monitor 23. This will generate relevant plots and data for Fig. 3G-bottom, which is the sleep timeseries for females.

#### 11.3. Replicating Fig. 4

1. Click the "…" button as shown in Fig. 1A-i and navigate to the folder containing the data *Set-1*.
2. Select all 32 channels of Monitor 23 (This is *Canton-S*, under LD12:12).
3. Set Exp. Start Date to 20-Dec-2020, Exp. Start Time to 15:01, Period Length to 24, Zeitgeber On to 6, and Hours of Zeitgeber to 12.

   ***Note:*** *Because peak detection requires the peaks of activity to be centered (and not at the edges of the cycle), we set Exp. Start Time to 15:01, which is six hours before lights-ON (21:00). This is why Zeitgeber On is set to 6 and because these flies are under LD12:12, the Hours of Zeitgeber is set to 12.*

4. Check the "Include Phase Analysis" button (Fig. 1B), and uncheck the "Include Activity/Sleep Analysis" button (Fig. 1A-iii).
5. Use default settings for phase picking and set Days to 1,2,3,4,5,6,7.
6. Type an output file name and navigate to any desired output folder location.
7. Click the "Averaged Activity Analysis" button (Fig. 1-A-v).
8. All relevant figures and data for Figs. 4A-left and 4B and C should have been generated.
9. Repeat the same steps as above but choose different monitors (to get all genotypes and photoperiods) and adjust the Exp. Start Time, Zeitgeber On and Hours of Zeitgeber depending on the photoperiod condition.

   ***Note:*** *Owing to the requirement of peaks to be centered, while analysing LD16:08 data, because lights-ON is at 21:00, the Exp. Start Time must be set to 17:01, the Zeitgeber On must be set to 4 and Hours of Zeitgeber must be set to 16.*

#### 11.4. Replicating Fig. 5

1. Click the "…" button as shown in Fig. 1A-i and navigate to the folder containing the data *Set-1*.
2. Select all 32 channels of Monitor 47 ($w^{1118}$).
3. Set Exp. Start Date to 25-Jun-2020, Exp. Start Time to 5:01, Period Length to 24, Zeitgeber On to 6, and Hours of Zeitgeber to 12.

   ***Note:*** *Because peak detection requires the peaks of activity to be centered (and not at the edges of the cycle), we set Exp. Start Time to 5:01, which is six hours before lights-ON (11:00). This is why Zeitgeber On is set to 6.*

4. Check the "Include Phase Analysis" button (Fig. 1B), and uncheck the "Include Activity/Sleep Analysis" button (Fig. 1A-iii).
5. Use default settings for phase picking and set Days to 1,2,3,4,5,6,7.
6. Type an output file name and navigate to any desired output folder location.
7. Click the "Averaged Activity Analysis" button (Fig. 1-A-v).
8. All relevant figures and data for Fig. 5A-top should have been generated.
9. Repeat the same steps as above but choose the monitor corresponding to the other genotype ($pdf^{01}$) to get all the remaining relevant figures and data for Fig. 5.

#### 11.5. Replicating Fig. 6

1. Click the "…" button as shown in Fig. 1A-i and navigate to the folder containing the data *Set-1*.
2. Select all 32 channels of Monitor 23 (*Canton-S*).
3. Set Exp. Start Date to 20-Dec-2020, Exp. Start Time to 15:01, Period Length to 24, Zeitgeber On to 6, and Hours of Zeitgeber to 12.

   ***Note:*** *Because peak detection requires the peaks of activity to be centered (and not at the edges of the cycle), we set Exp. Start Time to 15:01, which is six hours before lights-ON (21:00). This is why Zeitgeber On is set to 6.*

4. Check the "Include Latency or Anticipation Analysis" button (Fig. 1C), and uncheck the "Include Activity/Sleep Analysis" button (Fig. 1A-iii).
5. Use default settings for latency/anticipation analysis and set Days to 1,2,3,4,5,6,7.
6. Type an output file name and navigate to any desired output folder location.
7. Click the "Averaged Activity Analysis" button (Fig. 1-A-v).
8. All relevant figures and data for Fig. 6 (for *Canton-S*) should have been generated.
9. Repeat the same steps as above but choose all the channels from Monitor 56 ($per^{01}$) to get all the remaining relevant figures and data for Fig. 6.

### Section 12. Troubleshooting

#### 12.1. Troubleshooting saving and installing standalone installer on Mac OS

Complete one of the following steps before downloading the PHASE Installer on MacOS:

a. Allow downloads from unknown.
b. Open "System Preferences" by pressing the Apple logo on the top left of the screen.
c. In "Security & Privacy", under general, check the box to allow apps to be downloaded from "Anywhere".
d. If this option is hidden on MacOS version, open the "Terminal" and type this line of code: sudo spctl –master-disable. Click the return button. Enter the administrator password.
e. If unsuccessful with step (d), first download the program and then reopen the "System Preferences" window. At the bottom of the window will be the option for "Open Anyways".

#### 12.2. Occasional Excel errors while saving PHASE output

This may be due to running excel processes.

a. Go to Ctrl+Alt+Del and start Task Manager.
b. End all EXCEL processes
c. Try running PHASE again.

### References

Hendricks J, Finn S, Panckeri K, Chavkin J, Williams J, Sehgal A and Pack A (2000) Rest in Drosophila is a sleep-like state. Neuron 25:129–138. doi: 10.1016/S0896-6273(00)80877-6.

Huber R, Hill S, Holladay C, Biesiadecki M, Tononi G and Cirelli C (2004) Sleep homeostasis in Drosophila melanogaster. Sleep 27:628–639. doi: 10.1093/SLEEP/27.4.628.

Savitzky Abraham and Golay MJE (1964) Smoothing and Differentiation of Data by Simplified Least Squares Procedures. Analytical Chemistry 36:1627–1639. doi: 10.1021/AC60214A047.

Schafer RW (2011) What is a savitzky-golay filter? IEEE Signal Processing Magazine 28:111–117. doi: 10.1109/MSP.2011.941097.
